# Temporal and spatial domain-specific transcriptomic analysis of a vital reproductive meristem in *Arabidopsis thaliana*

**DOI:** 10.1101/032128

**Authors:** Gonzalo. H. Villarino, Miguel Flores-Vergara, Qiwen Hu, Bhupinder Sehra, Linda Robles, Javier Brumos, Anna Stepanova, Silvia Manrique, Lucia Colombo, Eva Sundberg, Steffen Heber, Robert G. Franks

## Abstract

**Background:** Plant meristems are analogous to animal stem cell niches as they maintain a pool of undifferentiated cells that divide and differentiate to give rise to organs. The carpel margin meristem is a vital, multi-potent structure located in the medial domain of the *Arabidopsis thaliana* gynoecium, the female floral reproductive organ. The carpel margin meristem generates ovules that upon fertilization become seeds. The molecular mechanisms that specify this meristematic region and regulate its organogenic potential are poorly understood. Here, we present an analysis of the transcriptional profile of the medial domain of the Arabidopsis gynoecium highlighting the developmental stages that immediately proceed ovule initiation, the earliest stages of seed development.

**Results:** Using a floral synchronization system and a *SHATTERPROOF2* domain-specific reporter, paired with fluorescence-activated cell sorting and RNA sequencing, we assayed the transcriptome of the gynoecial medial domain with temporal and spatial precision. This analysis reveals a set of genes that are differentially expressed within the *SHATTERPROOF2* expression domain that marks portions of the developing medial domain. Many members of this gene set have been shown previously to function during the development of medial domain-derived structures, including the ovules, thus validating our approach. Other uncharacterized members including differentially expressed cis-natural antisense transcripts, are potential novel regulators of medial domain development. Members of the *REPRODUCTIVE MERISTEM (REM)* family of transcriptional regulators were enriched in the *SHATTERPROOF2-expressing* cell population including a previously unrecognized *REM* family member. Finally, the analysis transcriptional isoforms in the medial domain identified genes that may exhibit “isoform switching” behavior during gynoecial development.

**Conclusions:** This data set provides genome-wide transcriptional insight into the development of the gynoecial medial domain that contains the carpel margin meristem, a vital reproductive structure that gives rise to the ovules in *Arabidopsis thaliana*.

## Background

The seedpod of flowering plants develops from the gynoecium, the female reproductive structure of the flower [1]. The gynoecium generates the ovules (the precursors of the seeds) and develops into the edible fruit in many fruiting species. As an estimated two-thirds of the calories of humankind's’ diet are derived from gynoecia and seeds, the gynoecium is a globally vital structure [2], [3].

In the flowering plant *Arabidopsis thaliana*, the gynoecium is a morphologically complex, multi-organ structure with a diversity of tissues and cell types [1, 4, 5]. The mature gynoecium displays morphological and functional differentiation along apical-basal, medio-lateral and adaxial-abaxial (inner-outer) axes. Stigmatic and stylar tissue form at the apex of the gynoecium, where the pollen grains are received and germinate. The stigma and style also comprise the apical-most portion of the transmitting tract, a structure that allows the pollen tube cell and sperm cells to reach the internally-located female gametophytes [4, 6, 7] (Fig. 1a,b,c). Located basal to the stigmatic and stylar tissue is the ovary portion of the gynoecium.

Ovules form within the ovary from a meristematic structure termed the medial ridge or carpel margin meristem (CMM), located in medial portions of the gynoecium [5, 8, 9] (Fig. 1a,b,c). Plant meristems are analogous to animal stem cell niches as they maintain a set of undifferentiated cells that can divide and differentiate into numerous tissues and cell types [10]. Early during floral development, patterning events divide the gynoecial primordium into medial domain that contains the carpel margin meristem and lateral domains that will form the walls of the gynoecium [5]. These domains express different sets of transcriptional regulators from early developmental time points.

Many genes that play a role in the development of the CMM and in the generation of ovules from this structure have been previously analyzed [8]. However, due to the complexity of the developing gynoecium and the heterogeneity of the gynoecial tissues, the ability to analyze the transcriptomic signature of the developing CMM or even other specific developing gynoecial structural domains has been limited. Wynn *et al*. previously evaluated the transcriptional properties of the gynoecial medial domain using hand-dissected gynoecial samples from the *seuss aintegumenta (seu ant)* double mutants that display a loss of many medial-domain-derived structures including ovules [11]. They identified 210 genes displaying reduced expression in *seu ant* gynoecia from floral stages 8-10 (stages according to Smyth *et al*. [12]). Many of these genes were shown via *in situ* hybridization to be preferentially expressed in the developing medial domain of the wildtype gynoecium and several of these genes have been shown to function during the development of ovules from the medial domain [8]. It is, however, difficult with this approach to obtain samples from gynoecia younger that stage 8 and thus to assay the earliest gynoecial patterning events.

An alternative approach to investigate the transcriptional properties of specific cellular populations utilizes Fluorescence-Activated Cell Sorting (FACS) of protoplasted cells to isolate specific-cell populations based on patterns of gene expression. This approach has been successfully applied to the Arabidopsis Shoot Apical Meristem (SAM) [13, 14] and roots [15, 16], [17], [18] as well as to developing cell lineages within the Arabidopsis leaf epidermis [19].

Here, we developed a novel FACS-based system for the transcriptomic analysis of a specific cellular population from the developing gynoecium, specifically the population of cells expressing the transcriptional regulator *SHATTERPROOF2 (SHP2). SHP2* encodes a MADS-domain transcription factor that is expressed early within the developing CMM and thus functions as a marker for the meristematic population of cells that generate the transmitting tract and ovules [20-23]. In order to focus our analysis on early stages of gynoecium development during which key patterning events occur, we generated a SHP2-domain-specific reporter in a genetic background that allowed the synchronization of floral development. FACS-based protoplast sorting procedures, coupled with RNA sequencing, provided a unique temporal and spatial precision to assay the transcriptional signature of the gynoecial SHP2-expression domain.

Our system provides the ability to isolate a large numbers of cells from a temporally- and spatially-restricted gynoecial domain. We apply this system to investigate the transcriptomic signature of the medial domain of the gynoecium at the developmental stages when key patterning events and ovule initiation occur. Our analysis reveals many genes that are expressed preferentially within the developing medial portions of the gynoecium including members of the *REPRODUCTIVE MERISTEM (REM)* family of transcriptional regulators [24, 25]. We also take advantage of strand-specific RNA sequencing technology to find coding protein genes and non-coding RNAs (ncRNAs) as well as to examine isoforms and naturally occurring antisense transcripts that are preferentially expressed in the medial domain.

This work complements and extends previous analyses of medial domain development and generates a list of potential novel regulators of medial domain development that are strong candidates for future functional analyses. Furthermore, global analyses of the transcriptomic dataset indicate a similarity of the pSHP2-expressing cell population to previously characterized meristematic domains, further supporting the meristematic nature of this gynoecial tissue.

## Results and Discussion

### FACS-based protoplast sorting allows the collection of the *SHP2-expressing* cell population from a temporally restricted inflorescence sample

The transcriptional regulator *SHP2* is preferentially expressed in the medial domain of the gynoecium and in a subset of the medial-domain derived tissues [21-23] (Fig. 1d,e). *SHP2* plays an important role in the development of the medial domain and in the specification of ovule identity [22, 26-29]. To better characterize the molecular mechanisms of the medial domain and ovule development, we sought to identify transcripts that are differentially expressed within the medial domain of the *Arabidopsis thaliana* gynoecium relative to the rest of the inflorescence. To enable this, we generated a transgenic line containing a two-component reporter system, in which a *pUAS-3xYPET* reporter was driven by a *pSHP2-GAL4* driver construct (Methods). Throughout this manuscript we refer to this two-component reporter as *pSHP2-YFP*.

**Figure 1.**
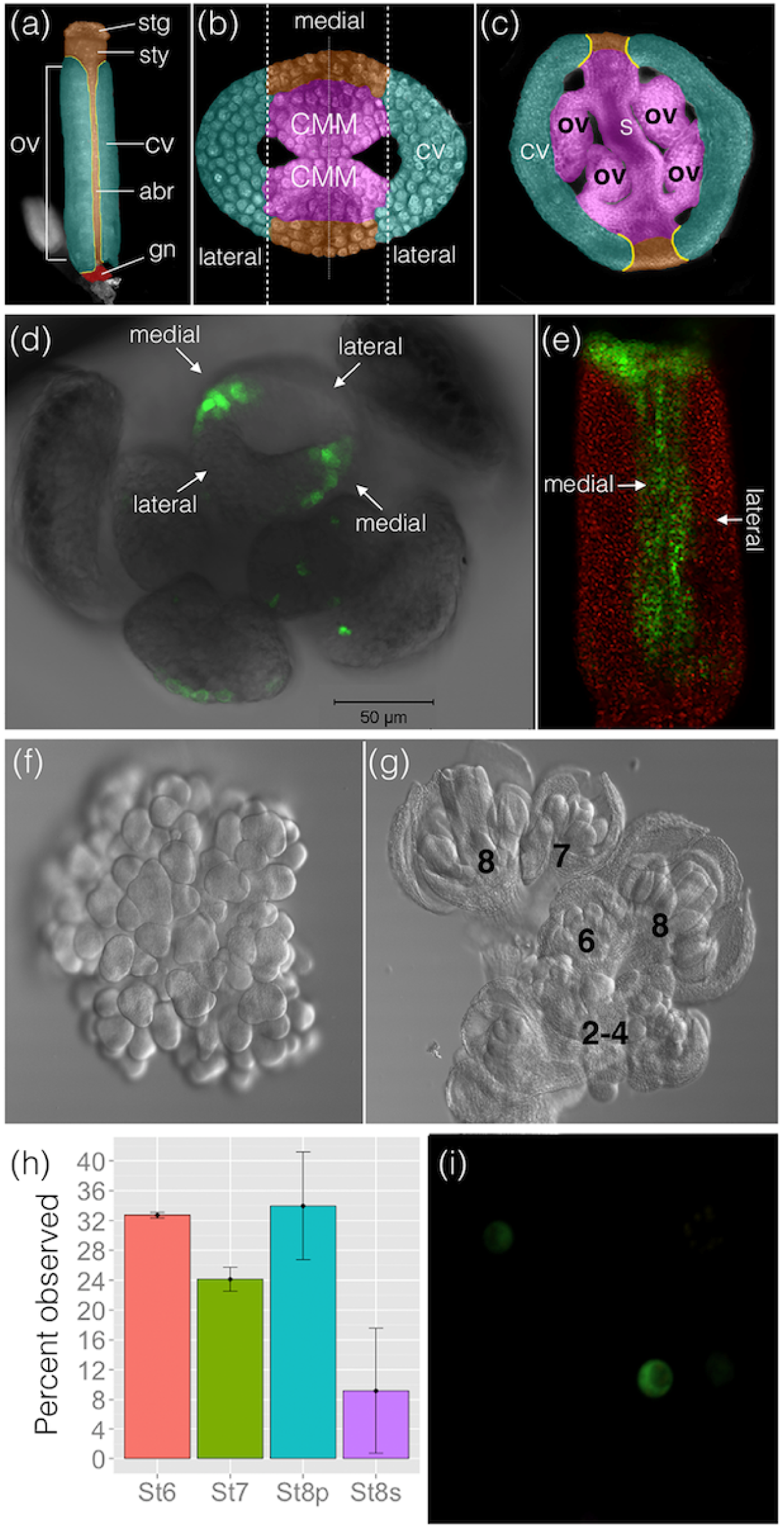
A system for the collection of temporally- and spatially-restricted cell populations from the *Arabidopsis thaliana* gynoecium. **a** Microscopic image of a mature wild type Arabidopsis gynoecium. The stigma (stg), style (sty), carpel valve (cv), abaxial replum (abr), gynophore (gn), and ovary (ovy) are false colored. **b** False-colored confocal cross section of a stage-8 gynoecium. Medial and lateral domains of the Arabidopsis gynoecium are indicated. The carpel margin meristem/medial ridge (CMM) is false colored pink. **c** False-colored stage-11 cross-section. Ovules (ov), septum (s) and carpel valves (cv) are indicated. **d** Confocal microscope image of the *pSHP2-YFP* two-component reporter in the *ap1; cal; pAP1::AP1:GR* background. YFP expression from the *pSHP2-YFP* reporter is chiefly confined to the medial domain of the gynoecium at late stage 7/early stage 8, although weak, non-medial domain expression can be detected in portions of the stamens. Sepals (se) and stamens (st) are labeled. **e** Z-stack composite 3D projection image of a gynoecium isolated from the flower at mid-stage 8. YFP expression from the *pSHP2-YFP* reporter is detected in the medial domain and at the apex of the gynoecium. **f** Chloral hydrate image of an inflorescence of an *ap1; cal; pAP1::AP1:GR* plant after mock treatment. Inflorescence-like meristems do not transition to floral meristems. **g** Chloral hydrate image of an inflorescence of an *ap1; cal; pAP1::AP1:GR* plant 125 hours after spray application of Dexamethasone synthetic hormone (Dex). Samples were enriched for stages 6-8. **h** Percentage of flowers at a given stage from inflorescences used for FACS-sorting. Stages 6, 7, 8p (pre-ovules) and 8s (post-ovules) are indicated in the X-axis as St6, St7, St8p, St8s, respectively. Stage 8p is before any visible morphological manifestation of ovule primordia upon observation under DIC microscopy. Stage 8s ovule primordia were observed and were at ovule stage 1-I or 1-II according the Schneitz *et al*. [125]. i Confocal microscopy of YFP fluorescence of protoplasted cells after FACS. Panels **a**, **b** and **c** are adapted from Azhakanandam *et al*. [43] (with permission).

To better understand the early specification of medial and lateral gynoecial domains and in the earliest stages of ovule primordium initiation, we focused our transcriptomic analysis on floral stages 6-8, when these key developmental events occur [5]. In order to increase our ability to collect a large number of *pSHP2-YFP-expressing* cells from this specific bracket of developmental stages, we crossed the *pSHP2-YFP* reporter into an *ap1 cal-based* floral synchronization system that allows the collection of large numbers of semi-synchronized flowers at roughly the same developmental stage [30, 31]. The expression of the *pSHP2-YFP* reporter in the floral synchronization system was largely similar to that observed in wild-type inflorescences [20, 22, 23], and was confined chiefly to the medial domain and medial domain-derived tissues (Fig. 1d,e). Some expression was observed in non-medial domain tissues. The most apparent of this was expression in the apex of the developing gynoecium where both medial and lateral domains express the *pSHP2-YFP* reporter. Additionally, expression could be observed in a small number of cells within the stamens (Fig. 1d) and occasionally in the edges of sepals that appeared to have undergone a homeotic transformation toward a carpelloid fate (data not shown). Thus, the vast majority of the *pSHP-YFP* reporter expression reflected the endogenous *pSHP2* expression domain (in the medial and apical portions of the gynoecium). A minority of the expression outside of the gynoecium may reflect ectopic expression of the reporter due to genetic background or transgene insertion site or limitations of the regulatory sequences used in the *pSHP-YFP* reporter construct.

Microscopic examination of our semi-synchronized inflorescence samples indicated that flowers ranged between floral stages 1 and early stage 8, with a strong enrichment for floral stages 6 through early 8 (Fig. 1g,h). Flowers that had developed beyond late stage 8 were not detected in our samples. Thus, our biological sample is strongly enriched for transcripts that are expressed during early patterning of the gynoecium and the earliest stages of ovule development (initiation) and does not include later floral developmental stages where *SHP2* is expressed in stigma, style and valve margin tissues. Additionally, as the initial expression of the *pSHP-YFP* reporter is detected at late stage 5 or early stage 6 [23], we expect that the population of YFP-expressing protoplasts derived from this material will be highly enriched with cells from the stage 6-8 medial domain. FACS-sorting of protoplasts derived from these inflorescences yielded three populations of sorted cells (collected in biological quadruplicate): “YFP-positive”, “YFP-negative” and “all-sorted” (Additional file 1: Figure S1). The “all-sorted” sample included all protoplasts recovered (regardless of YFP expression) after sorting gates were applied to remove debris and broken cells (Methods). We additionally collected (also in biological quadruplicate) “non-sorted” samples from entire non-protoplasted inflorescences to measure the abundance of transcripts in the biological starting material before protoplast generation and FACS-sorting. In order to evaluate the purity of the YFP-positive protoplasts during a preliminary FACS run, YFP-positive cells were resorted. Ninety six percent of the YFP-positive cells were found to resort into the YFP-positive gate, indicating a high degree of enrichment and purity in the YFP-positive sample (Additional file 1: Figure S1). Confocal microscopy (Fig. 1i) also revealed an enriched population of intact YFP-positive protoplasts after FACS.

We used real time PCR (qRT-PCR) to estimate the degree of enrichment of the endogenous *SHP2* and *NGATHA1 (NGA1)* transcripts in RNA samples derived from the YFP-positive and YFP-negative samples. *NGA1* is expressed in the adaxial portions of the gynoecium starting at stage 7 in a domain that partially overlaps with the *SHP2* expression domain [32, 33] and thus provides an additional benchmark to estimate the enrichment of medial domain-expressed transcripts. The normalized level of the *SHP2* transcript was ~30 fold higher in the YFP-positive samples relative to the YFP-negative samples *(p* <0.001) while the *NGA1* transcript was ~4 fold higher in the YFP-positive sample *(p* <0.05). The difference in the levels of the *TUBLIN6* was not found to be statistically significant *(p* = 0.4) between the YFP-positive and YFP-negative samples (Additional file 1: Figure S2).

### Transcriptomic analysis of the gynoecial *SHP2* expression domain and identification of candidate regulators of gynoecial medial domain development

To investigate the transcriptomic profile of the gynoecial *SHP2* expression domain, we performed high-throughput RNA-sequencing from the collected protoplasts and non-protoplasted inflorescences samples. We expect that the identification of differentially expressed genes (DEGs) between the YFP-positive and YFP-negative samples (referred to as “YFP-positive/YFP-negative” or “YFP+/-”) will provide insight into the set of transcripts differentially expressed in the gynoecial medial domain relative to the rest of the inflorescence. Additionally, DEGs identified in the all-sorted and non-sorted comparison (referred to as “all-sorted/non-sorted”) are expected to reveal transcripts that are differentially represented as a result of the protoplasting/FACS-sorting protocol.

Two lanes of the HiSeq2500 Illumina sequencing platform yielded 320 million raw reads with an average of 20 million reads (MR) per library. Nearly 11 MR were filtered out after removing barcode-adapters and low quality sequences. The remaining 306 MR were aligned against the *Arabidopsis thaliana* TAIR10 reference genome [34] with more than 90% of them successfully mapping to the genome sequence. Among the mapped reads, 244 MR mapped uniquely to only one location and were used for subsequent analyses. A detailed breakdown is shown in Additional file 2: Table S1.

We used three different programs to determine expressed and differentially expressed protein coding genes in our dataset: Cufflinks [35], edgeR [36] and DESeq2 [37] (Methods) (Non-protein coding gene models were considered separately and are presented below). Here, the term “differentially expressed gene (DEG)” is used to indicate a gene whose steady-state transcript level differs significantly at a false discovery rate (FDR) of <0.001 and shows a fold change of four or more between the two compared RNA samples. To identify potential regulators of gynoecial medial domain development, a ‘stringent’ criteria was used to select a subset of the YFP+/-DEGs for downstream analysis. For a gene to be selected from the YFP+/-comparison, we required that the transcript is identified as differentially expressed by all three independent software packages (Fig. 2b). Alternatively, to identify DEGs in response to the protoplasting/FACS-sorting procedure, a ‘less stringent’ criterion was used. Transcripts in the union set of all the non-sorted/all-sorted DEGs were considered to be potential protoplast-induced genes even if they were identified by only one software program (Fig. 2a). Only 48 transcripts were found in common between the YFP+/-DEGs and the all-sorted/non-sorted DEGs (Fig. 2c), indicating a high degree of specificity in the DEGs identified in each comparison. We then removed these 48 transcripts from our analysis to eliminate any that might be differentially expressed as a result of the protoplast generation or FACS-sorting procedures, leaving 363 “cleaned” protein coding DEGs (Fig. 2c). The expression profiles of these 363 YFP+/-DEGs, including data from the all-sorted and non-sorted samples, are represented in a heatmap (Additional file 1: Figure S3). This gene set includes 95 DEGs whose transcript levels were higher in the YFP-positive samples (“enriched”) and 268 DEGs whose transcript levels were lower (“depleted”) in the YFP-positive samples, relative to the YFP-negative samples (Additional file 3: Table S2).

**Figure 2.**
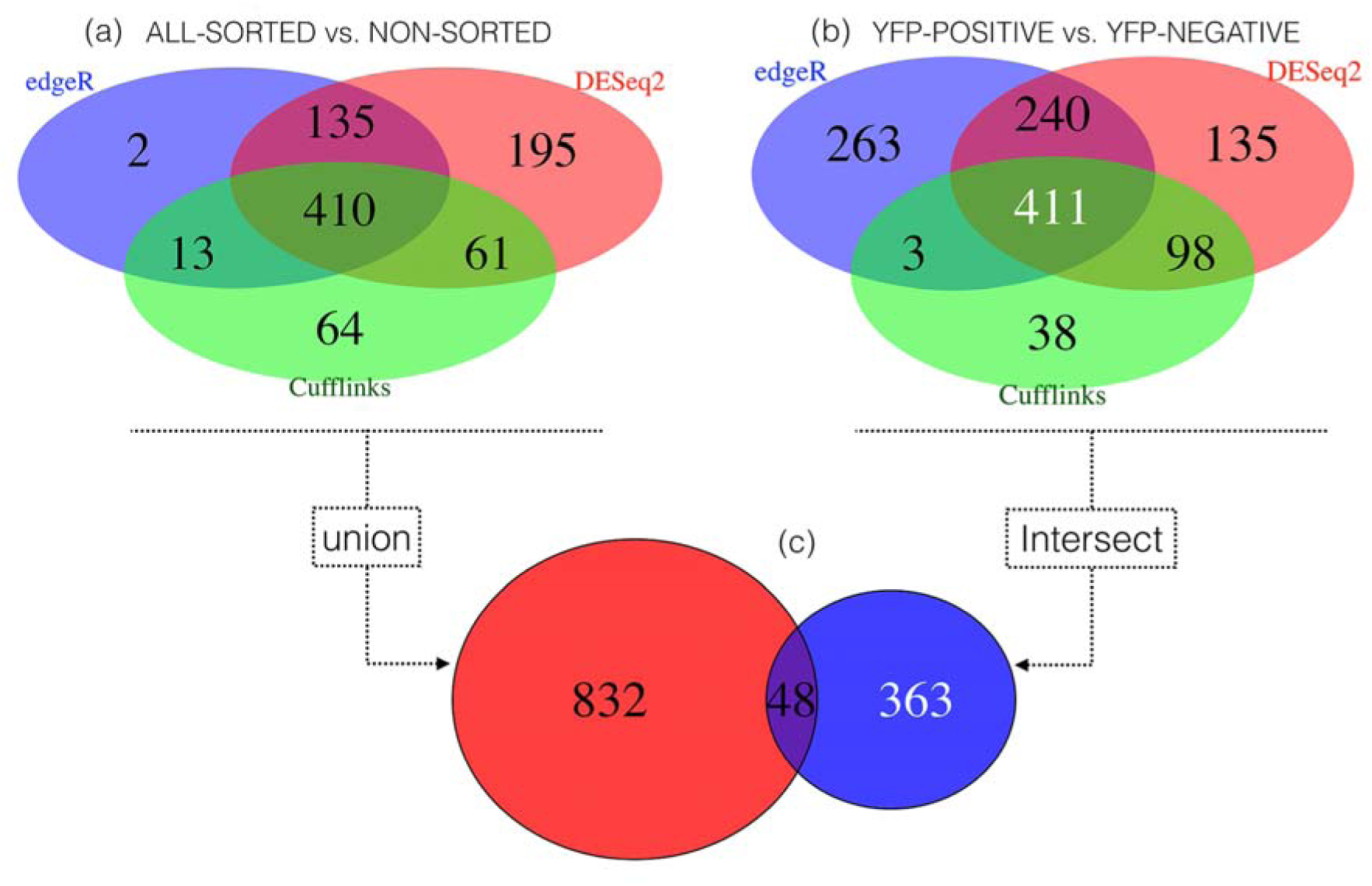
Venn diagram of differentially expressed genes (DEGs) using Cufflinks, edgeR and DESeq2 (FDR<0.001, Fold Change >4). **a** Venn showing DEGs identified between the all-sorted/non-sorted samples with the 3 programs used for differential expression analysis of RNA-seq expression profiles. **b** Venn showing DEGs between YFP+/-samples identified in the 3 programs. **c** Intersection of the DEGs (48) from both datasets (a and b). DEGs (363), after removing DEGs induced by the protoplsting/FACS-sorting stress, were used for downstream analysis.

For the 95 DEGs that were enriched in the YFP-positive sample (at a fold change > 4), we expected many to be preferentially expressed in the medial portions of the gynoecium at floral stages 6-8. To test this, we examined the literature to determine the expression patterns of members of this gene set. From the top 15 of the 95 YFP-positive enriched DEGs (ranked by fold change), five have previously been reported to be preferentially expressed in the gynoecial medial domain via *in situ* or reporter gene analysis [i.e. *HECATE1 (HEC1), HEC2, SHP1, SHP2* and *STYLISH1 (STY1)]* [20-22, 38, 39] and three others are previously described as enriched in medial domain-derived tissues in published transcriptomic datasets (i.e., AT1G66950, AT5G14180, and AT1G03720) [40, 41] (Table 1). An additional gene from this list, *CRABS CLAW (CRC)*, has been shown via *in situ* hybridization to be expressed in portions of the medial gynoecial domain, as well as non-medial portions of the gynoecium [42, 43]. The expression pattern of the remaining six genes from this gene list have not yet been assayed in the gynoecium. Thus, as predicted, the set of 95 genes enriched in the YFP-positive sample is enriched for genes that are preferentially expressed in the gynoecial medial domain.

**Table 1.** Top 15 most differentially expressed genes from the YFP+/-comparison (FDR <0.01), enriched in the YFP-positive sample ranked by fold change (FC). Arabidopsis gene ID is shown in the 1^st^ column, gene name (TAIR10 annotation) is shown in the 2^nd^ column and the 3^rd^ column reference available reporter lines and/or *in situ* hybridization for the top 15 DEGs in this study. Reads Per Kilobase of transcript per Million mapped reads (RPKM) values are indicated for each sample (YFP_NEG = YFP-negative, YFP_POS= YFP-positive) and each biological replicates (B1, B2, B3, B4).

Published functional analyses of *HEC1, HEC2, SHP1, SHP2* and *STY1* indicate that these genes function during the development of the medial domain or medial domain-derived tissues [22, 28, 29, 38, 39]. Many other genes in the set of 95 DEGs enriched in the YFP-positive sample have been previously shown to play a role in medial domain development (e.g. *NGA* family members [32, 33], SPT [44], and *CUC2* [45]). Other genes within this list are interesting candidates for future functional studies. This includes members of the *REM* family of transcriptional regulators [24, 25], several auxin synthesis or signaling-related genes such as *LIKE AUXIN RESISTANT 1 (LAX1)* (AT5G01240) [46] and *YUCCA4 (YUC4)* (AT5G11320) [47], as well as transcription factors regulating other developmental processes such as *MATERNAL EFFECT EMBRYO ARREST 3 (MEE3)* (AT2G21650) [48] and *GLABROUS 3* (AT5G41315) [49].

It is important to note that the 48 DEGs that were identified in both the YFP+/- and all-sorted/non-sorted comparisons (Fig. 2c) should not be discounted as potential medial domain regulators. These genes may be both preferentially expressed in the YFP-positive cell population as well as induced in response to the protoplasting procedure (Additional file 3: Table S2). Indeed, some of these genes, including the transcription factors *HECATE3* and *BR-ENHANCED EXPRESSION1* (BEE1), have been reported to be preferentially expressed in medial domain-derived tissues and to function in gynoecium development [38], [50]. However, we chose to use the “cleaned” set of 363 YFP+/-DEGs for downstream analyses in order to reduce the likelihood of the inclusion of genes whose expression was altered significantly by the protoplasting process.

### *REPRODUCTIVE MERISTEM* family members are differentially expressed in the SHP2-expression domain

In order to look for enriched categories of transcription factors within the set of “cleaned” 363 YFP+/-DEGs (Fig. 2c), we used the online Transcription Factor Enrichment Calculator tool [51]. Members of the *ABI3/VP1* transcription factor family that includes the *REM* family TFs and *NGA* family TFs were found to be statistically over-represented (Additional file 8: Table S7) (corrected *p* <9.97E-06). The *REMs* belong to the plant-specific B3 superfamily of transcription factors and expression of many REM family members is observed in meristematic tissues such as the inflorescence meristem, floral meristem and the CMM [11, 24, 25, 52-54]. The numerical designations used to describe the *REM* family members in this manuscript are taken from Romanel *et al*. [24]. In our study, six *REM* members were amongst the 363 statistically significant YFP+/- DEGs; five were found to have enriched expression in the YFP-positive sample, while one, *REM25 (AT5G09780)*, was ~4 fold less abundant in the YFP-positive sample. *REM13* (At3g46770) transcript level is enriched ~12 fold in the *pSHP2-YFP* expressing cells. *REM13* was previously predicted to be preferentially expressed in the inner integument, ovule primordia and medial domain based on transcriptomic data [40]. We employed *in situ* hybridization to assay the expression pattern of the *REM13* transcript during gynoecial development (Figure 3). Using a *REM13* antisense probe, we detected signal in the medial portions of the gynoecium corresponding to the carpel margin meristem as early as stage 7. Expression was also observed in the initiating ovule primordia in stage 8 gynoecia and then continued to be detected in portions of the ovules at later developmental stages.

**Figure 3.**
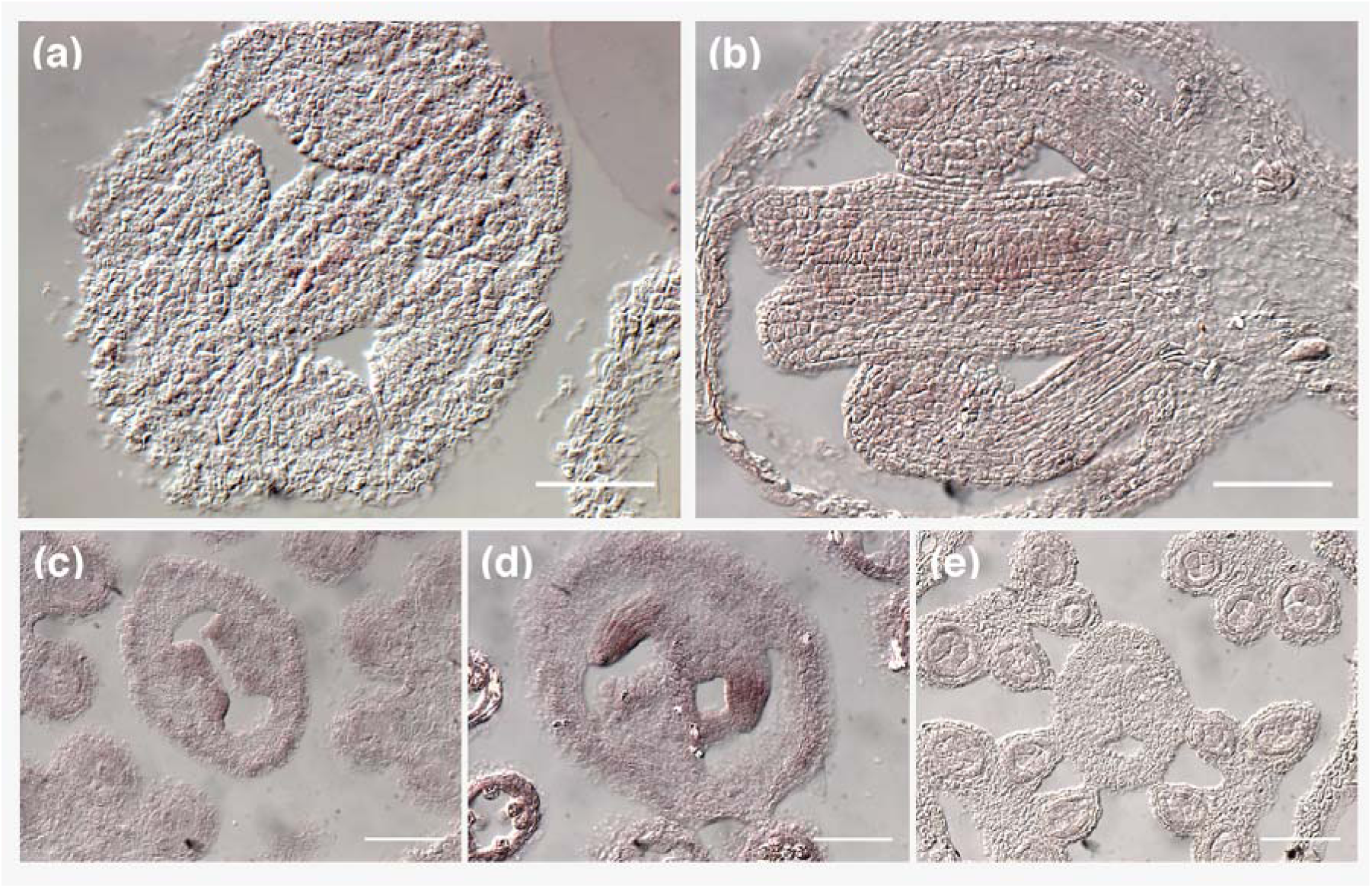
Candidate medial domain regulator *REM13* (At3g46770) is expressed within the medial gynoecial domain and developing ovules. Results from an RNA *in situ* hybridization with *REM13* probe. **a-d** antisense probe. **e** sense strand probe. **a** Hybridization signal is detected in the carpel margin meristem (adaxial portions of the medial gynoecial domain) in the stage 7 longitudinal section. **b, c** and **d** In transverse gynoecial sections *REM13* expression is detected in the ovule primordia; stage 7 (panel b) stage 8 (panel c) and stage 9 (panel d) gynoecia. **e** A stage 8 section hybridized with a *REM13* sense strand probe. (ov) - ovules, (cmm) - carpel margin meristem. Scale bars for each panel represent 5 microns.

*REM34/ATREM1* (At4g31610) [24, 52], *REM 36* (At4g31620)[53], and *VERDANDI (VDD/REM20)* [53, 55] also displayed enriched expression levels in the YFP-positive sample of ~8 fold, ~9 fold and ~6 fold, respectively. Published *in situ* hybridization patterns indicate enriched medial domain expression patterns for *REM34/ATREM1* and *VDD/REM20* [11, 52, 55]. Additionally, expression of At5g60142, a previously unnamed member of the *REM* family, is enriched ~11 fold in the YFP-positive sample (Additional file 4: Table S3). At5g60142 is an interesting candidate for functional studies that is located on chromosome V in tandem to *REM11* (At5g60140) and *REM12* (At5g60130) and shares a high degree of sequence similarity with these two genes, as well as *REM13* [24, 53]. We propose to designate At5g60142 as *REM46*.

### Gene Set Enrichment Analysis

To gain global insights into underlying biological mechanisms of medial domain development and function, Gene Set Enrichment Analysis (GSEA) was performed for the 95 YFP-positive enriched DEGs in the medial domain. This analysis identified 147 GO terms that were statistically overrepresented (*p* < 0.01), including “gynoecium development” (GO:0048467) and “flower development” (GO:0009908), “response to gibberellin” (GO:0009739) and “auxin homeostasis” (GO:0010252) (Fig. 4 and Additional file 7: Table S6). This GSEA analysis further suggests that the set of 95 genes enriched in the YFP-positive sample function as regulators of medial domain development.

**Figure 4.**
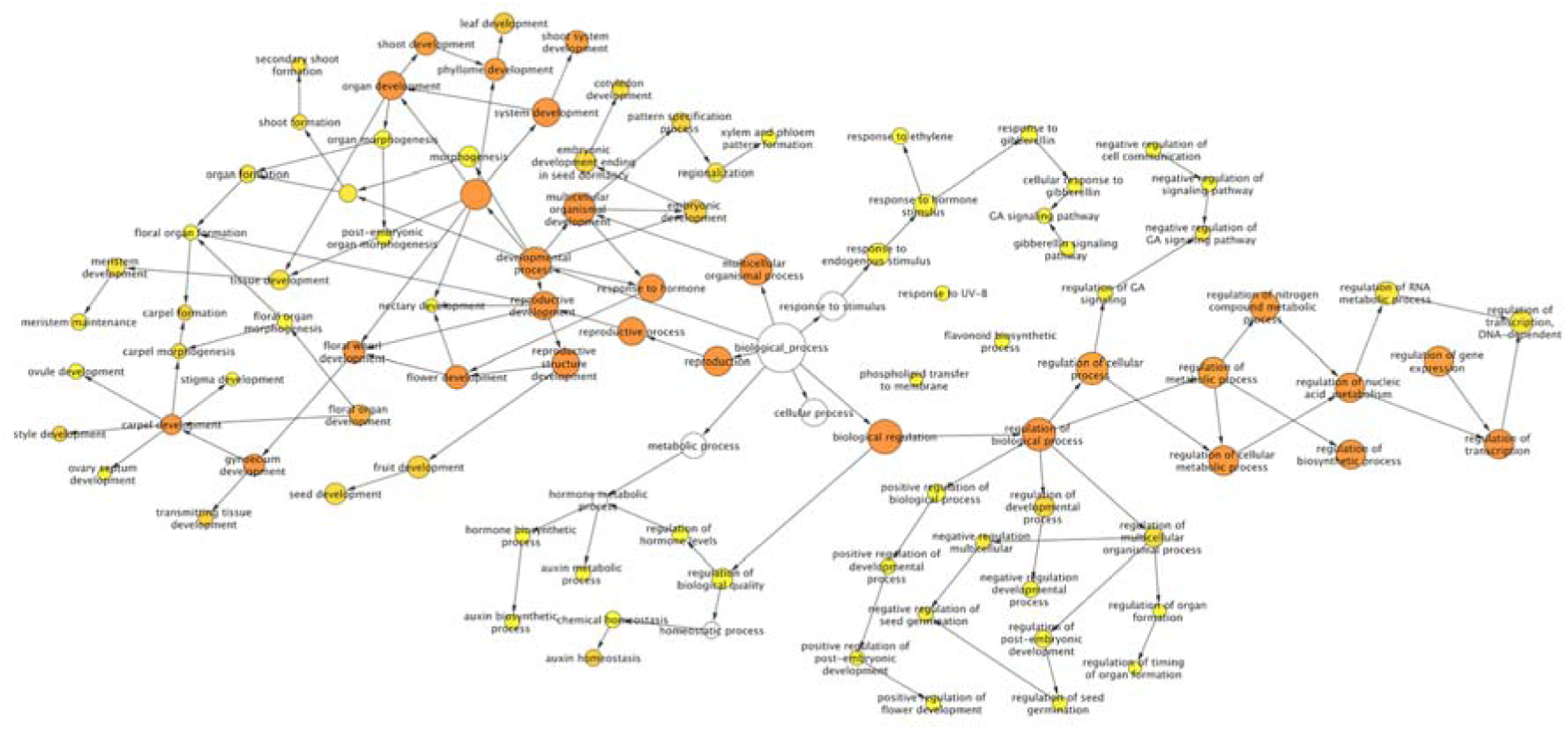
GO term overrepresentation of SHP2-domain enriched genes suggests a role for this set of genes in floral, gynoecial and ovule development. BiNGO/Cytoscape representation of overrepresented GO terms from the 95 YFP+/- DEGs displaying enriched expression in the YFP-positive samples. Edges represent the parent/child relationships of the GO terms [126], while color of the nodes indicates the degree of statistical significance *(p* <0.01) as reported by BiNGO [123]. To unclutter the figure, given the large number of significant GO terms, selected nodes and edges have been removed from this graphical representation.

In contrast, when performing GSEA with DEGs identified between the all-sorted/nonsorted samples, a different set of 304 overrepresented GO terms were identified, including “response to stress” (GO:0006950) and “response to wounding” (GO:0009611), suggesting that many of the genes identified as differentially expressed between the all-sorted/non-sorted samples reflect stress-induced changes in gene expression during protoplast/FACS-sorting.

### The transcriptomic signature of the SHP2-expressing cell population shares commonalities with transcriptional signatures of other meristematic samples

In order to gain insight into the characteristics of the 363 YFP+/- DEGs identified from the *SHP2* expression domain, we compared the expression profile of this set of genes across several different tissues. Using existing Arabidopsis RNA-seq transcriptomic datasets from whole flowers [56], aerial seedlings tissues (GEO accession: GSE54125), as well as from Laser Capture Microdissected (LCM) inflorescence meristems, floral meristems and stage-3 flowers [54], a Spearman rank correlation analysis was performed. In the sample-wise hierarchical clustering (Fig. 5a), the transcriptomic profiles from the SHP2-expressing (YFP-positive) sample clustered more closely with the meristematic samples, while the YFP-negative and all-sorted samples clustered more closely with the whole-flower and whole-seedling samples. This suggests that the expression signature of the YFP-positive sample is more similar to that of the floral and inflorescence meristems and young flowers, than it is to whole flowers or young vegetative seedlings (Fig. 5a).

**Figure 5.**
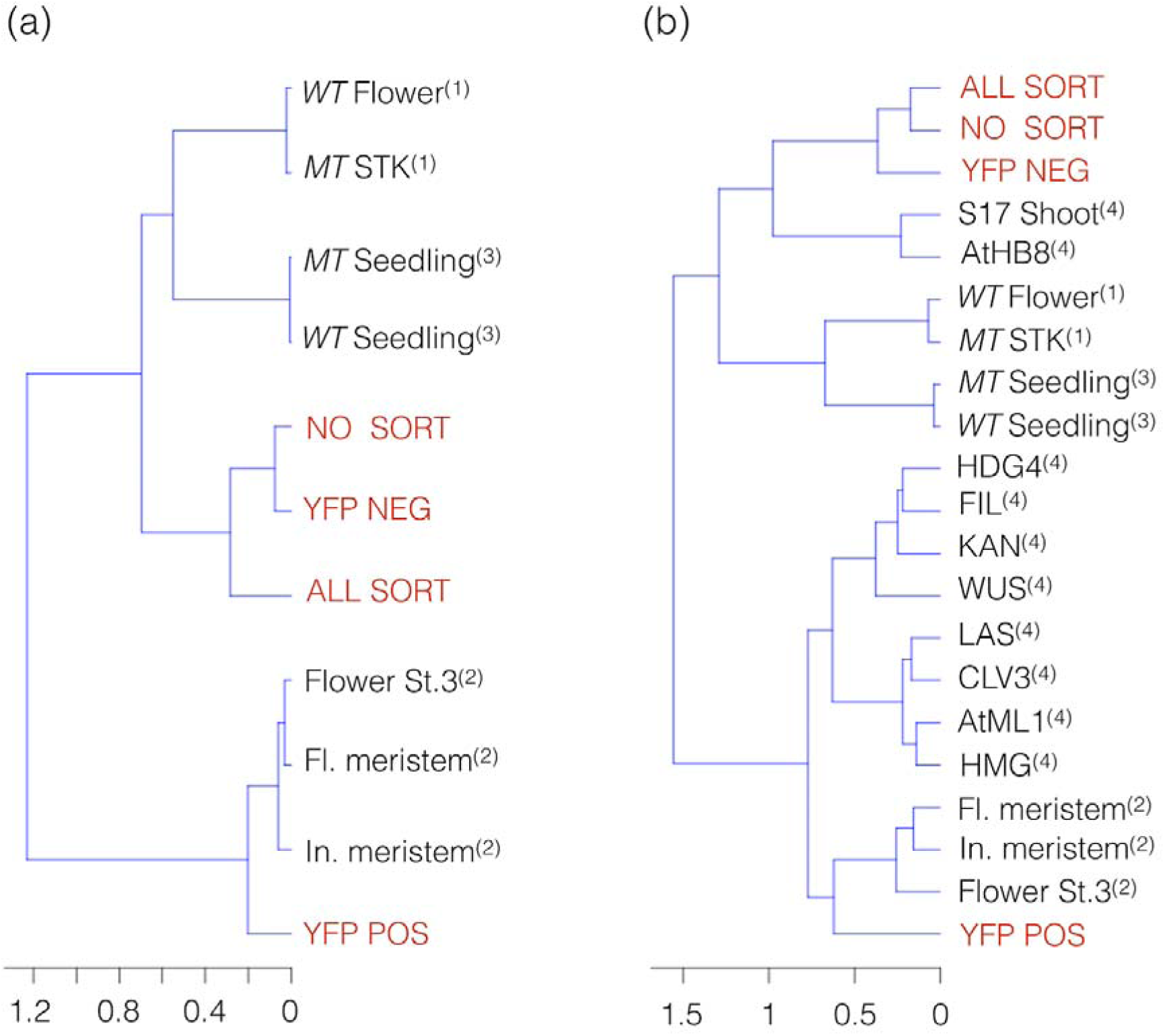
The transcriptomic signature of the *SHP2-expressing* domain is more similar to the transcriptomes of other meristematic samples than it is to whole flower. **a** Dendrogram based on hierarchical clustering using the Spearman rank correlation using RNA-seq (RPKM) expression values from flowers and other tissues. **b** Comparison of RNA-seq and affymetrix ATH1 arrays samples including transcriptomic data from whole flower, shoot apical meristem and seedling. WT = wild type, MT= mutants. Data from Mizzotti *et al*. [56]^(1)^, Mantegazza *et al*. [53]^(2)^, GEO accession: GSE54125^(3)^ and Yadav *et al*. [13, 14]^(4)^ were used for comparison. Samples corresponding to this study are color coded red in both dendrograms.

Further supporting the similarity of the SHP2-expressing domain to other meristematic samples, the expression levels of *GA20OX1* (AT4G25420) and *GA20OX2* (AT5G51810) were both significantly depleted in the YFP-positive sample, relative to the YFP-negative sample (Additional file 11: Table S10). *GA20OX1* and *GA20OX2* encode key biosynthetic enzymes of the plant hormone gibberillic acid (GA) [57]. Levels of expression of *GA20OX1* and *GA20OX2* are low in the shoot apical meristem (SAM) relative to expression in the juxtaposed young organ primordia and high levels of GA synthesis interfere with the maintenance of meristematic fate in the SAM [58, 59]. These data suggest that low levels of GA may also be associated with the meristematic nature of the carpel margin meristem. Although not discussed here, expression values of genes annotated with a role in ethylene signaling are found in Additional file 11: Table S10.

We additionally compared the medial domain transcriptional signature to datasets generated with the Affymetrix ATH1 array allowing comparisons to transcriptomic signatures of a variety of cell types including vascular and meristematic cell types from the Arabidopsis SAM isolated via FACS [13, 14]. When these additional samples are included, the hierarchical clustering dendrogram (Fig. 5b) shows the YFP-positive sample is more similar to the SAM cell-types, rather than to the vascular procambium (AtHB8) and phloem cell types (S17). This again suggests the meristematic character of the YFP-positive sample (Fig. 5b). One should be cautious, however, to interpret the results of this (or any) cross-platform (array/RNA-seq) comparison until validated crossplatform comparisons methods are available. To the best of our knowledge, there is no clear consensus in the literature of a standard cross-platform comparison practice [60], [61], [62], [63]. Indeed, many researchers have used both platforms (array/RNA-seq) in the same experiment comparing final results rather than finding a way to directly compare the two technologies [64], [65], [66], [63], [67]. Here, we employ a Spearman rank correlation as it is less sensitive than the Pearson correlation to strong outliers, makes no assumptions about data distribution, and does not inflate type I error rates. This approach fits well with the data in this work as samples do not cluster based on technology platforms but rather cluster based on the apparent cell-type similarities of gene RPKM (Reads Per Kilobase of transcript per Million mapped reads) expression levels.

### Transcriptomic analysis of the *SHP2* expression domain complements existing medial domain and CMM data sets

Wynn *et al*. previously carried out a related transcriptomic study and identified many genes that were shown via *in situ* hybridization to be preferentially expressed in the developing medial domain of the wild-type gynoecium [11]. When comparing the 95 enriched DEGs from our RNA-seq experiment (Additional file 3: Table S2 and Additional file 1: Figure S3) with a set of 210 medial domain enriched genes from Wynn *et al*., 23 genes were found in common (Table 3). The 24% overlap of these two gene sets is significantly higher than expected by chance (hypergeometric test; *p* = 3.15 × 10^−30^) [68]. Members of the *REM, HECATE* and *NGA* gene families, as well as several auxin-homeostasis-related genes were among the set of 23 genes identified in both experiments (Table 3).

Reyes-Olalde *et al*. recently performed a comprehensive literature survey of genes that function during CMM development [8]. They reported 86 protein-coding genes corresponding to transcription factors, hormonal pathways, transcriptional co-regulators, and others of widely diverse functions. While all 86 are expressed in our dataset, fifteen of these CMM developmental regulators are found within the set of 363 YFP-positive DEGs (hypergeometric test; *p* = 3.3 × 10^−13^) [68] (Fig. 6). The expression profiles of the 86 genes reported by Reyes-Olalde *et al*. within the medial domain-enriched dataset from this work, as well as within data from floral meristem enriched samples [54], is displayed in a heatmap in Figure 6 (RPKM values can be found in Additional file 10: Table S9).

**Figure 6.**
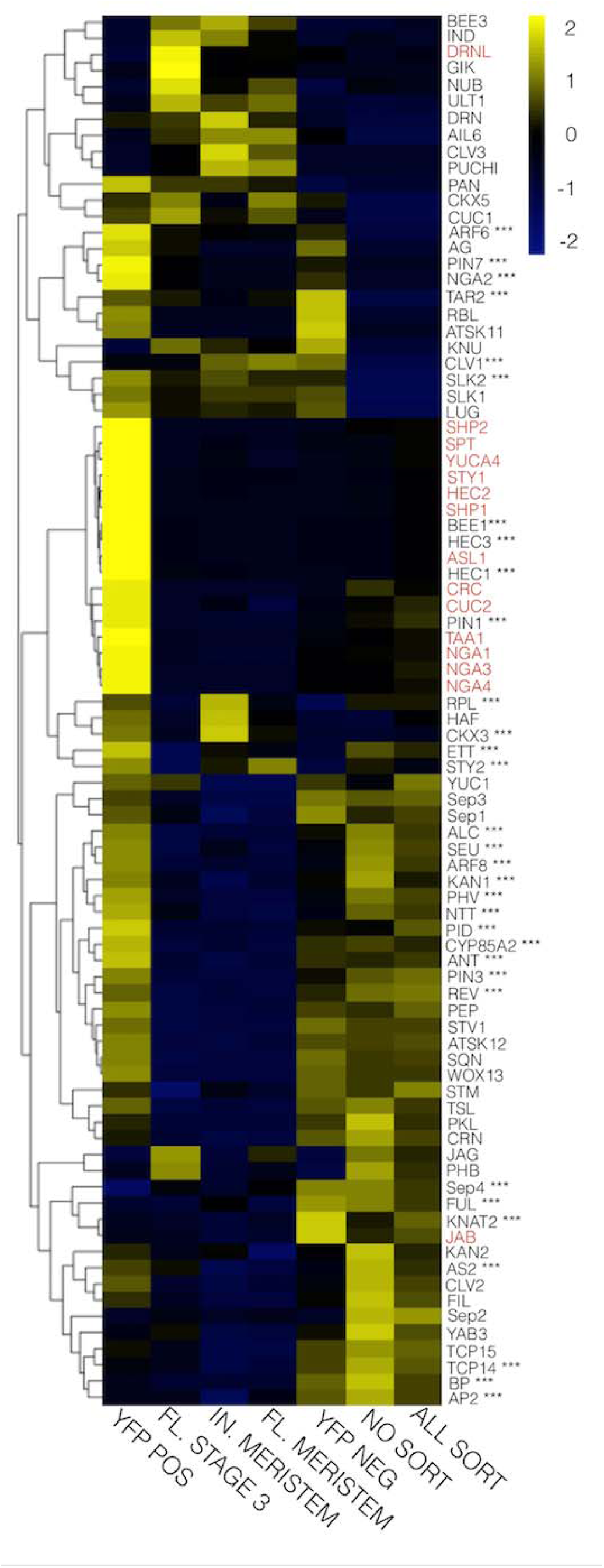
Heatmap representation of the expression profiles of previously identified regulators of Carpel Margin Meristem development. Expression profiles in Reads Per Kilobase of transcript per Million mapped reads (RPKM) of the 86 genes reported by Reyes-Olalde *et al*. [8] with functional role during CMM development. Transcriptional profiles from this study (YFP POS = YFP-positive, YFP NEG = YFP-negative, ALL SORT = all-sorted, and NO SORT = no-sorted) as well as Mantegazza *et al*. [54] corresponding to flower stage 3 (FL.STAGE 3), floral meristem (FL.MERISTEM) and inflorescence meristem (IN.MERISTEM) are included. Genes color-coded in red are those identified as DEGs between YFP-positive and YFP-negative samples (FC >4 and FDR <0.001) while genes that displayed a statistically significant expression level (FDR <0.01) between YFP-positive and YFP-negative (regardless of their fold change) are indicated with ^***^.

### Transcript isoforms in the *Arabidopsis* medial domain

One utility of transcriptome analysis through RNA-seq is the identification of novel alternative spliced transcripts, alternative transcription start sites (TSS), and instances of isoform switching [69]. To further characterize the transcriptome of the SHP2-expression domain at the isoform level, we first selected isoforms that showed a significant (α <0.01) change in their expression between YFP+/-samples using Cufflinks/Cuffdiff. For this analysis we did not apply a fold magnitude cutoff, thus capturing all isoforms with α <0.01. To avoid transcripts that were affected by the cell-sorting procedure, we removed all isoforms that showed a significant (α <0.01) expression level change between all-sorted/non-sorted samples. This resulted in 4555 YFP+/- differentially expressed isoforms (Additional file 9: Table S8). Within this set of isoforms differentially expressed between the YFP+/- samples, we sought to highlight multi-isoform genes that showed major changes in the relative frequency of individual isoforms between the YFP-positive and YFP-negative samples. To this end, we estimated the relative frequency of each isoform as a percentage of the total expression for the gene. Among the 4555 significantly differentially expressed isoforms, only 52 isoforms from multi-isoform genes displayed changes of 20% or more in their relative frequency. The major isoform (most highly expressed isoform) differed between YFP+/- samples for only 15 genes (Table 2). Remarkably, the transcriptional co-regulator SEU (At1g43850), previously implicated in medial domain development [70], [43], showed a significant increase of isoform At1g43850.1 in the YFP-positive samples, while its second isoform At1g43850.2 did not significantly change between samples. As a result, isoform 1 was the major (predominant) isoform in YFP-positive cells, and isoform 2 was the major (predominant) isoform in the other samples. The functional significance, if any, of this isoform switching is currently unknown.

**Table 2.** List of significant isoforms (α=0.01) between the YFP+/- samples and nonsignificant (a=0.01) between the all-sorted/non-sorted comparison. These genes were 20% or more enriched in the medial domain for a given isoform. TSS = transcriptional start site. Match between the Cufflinks transcripts and TAIR10 genome are indicated with class code ‘=’ for complete transcript match and ‘j’ for potentially novel isoform (fragment) [35].

**Table 3.** List of overlapping differentially expressed genes (DEGs) between the 95 DEGs from this study (FC > 4, FDR <0.001) and DEGs from the transcriptomic array data of Wynn *et al*. [11] derived from the *seuss aintegumenta (seu ant)* double mutant.

The regulation of gene expression through alternative promoter usage or use of alternative TSS is frequently observed in multicellular organisms [71]. Using the same pipeline and criteria we employed to select differentially expressed isoforms in the YFP+/- samples, we identified 93 isoforms that were differentially expressed as a result of the use of alternative promoter/transcriptional start sites (Additional file 9: Table S8). Interestingly, one such promoter/transcriptional start site switch was found for the

*REVERSIBLY GLYCOSYLATED POLYPEPTIDE 5 (RGP5)* gene (isoform). Members of the *RGP* family *(RGP1* and *RGP2)* involved in sugar metabolism are expressed in other Arabidopsis meristematic tissues, such as the root tip and the apical meristem of young seedlings [72]. In our work, the transcript level of *RGP5* isoform 2 (At5g16510.2) in the YFP positive sample is 61% higher relative to the level of this isoform in the YFP-negative sample, while the level of isoform 1 (At5g16510.1) is 75% lower (Fig. 7a and Additional file 9: Table S8).

**Figure 7.**
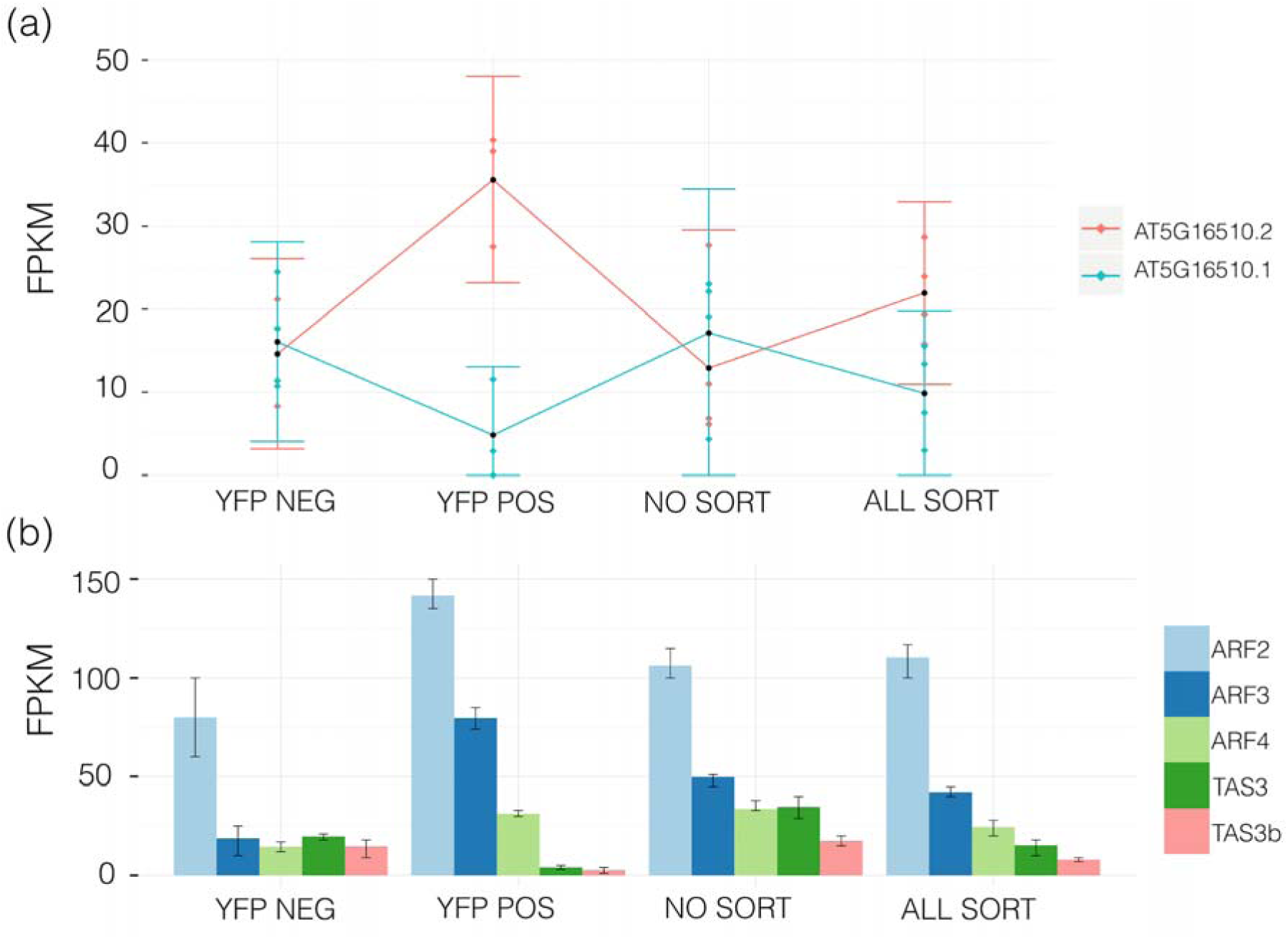
Differential expression of *REVERSIBLY GLYCOSYLATED POLYPEPTIDE 5 (RGP5)* isoforms as well as *TRANS-ACTING siRNA3 (TAS3)* and *AUXIN RESPONSE FACTOR* genes. **a** Promoter/transcriptional start site switch found for the *RGP5* gene (At5g16510). The isoform 2 (At5g16510.2) increases its expression in the YFP-positive domain while isoform 1 (At5g16510.1) of the same gene decreases its expression in the same domain. **b** Expression of the *AUXIN RESPONSE FACTORS (ARFs) (ARF2, ARF3, ARF4)* and *TAS3* transcripts. Expression levels of *ARF2, ARF3, ARF4* are significantly enriched in the YFP-positive sample at FDR <0.01. Expression levels of the *TRANS ACTING siRNA3 (TAS3)* genes At5g49615 and At3g17185, that negatively regulate the expression of *ARF2, ARF3*, and *ARF4* expression [85], are significantly reduced (FDR <0.01) in the YFP-positive sample.

### Auxin homeostasis and the development of the gynoecial medial domain

Auxins are a class of plant hormones that regulate growth and development [73, 74]. The most common plant auxin is Indole-3-Acetic Acid (IAA). The regulation of auxin homeostasis (including synthesis, response, transport, inactivation and degradation) plays an essential role in patterning the gynoecium and other lateral organs [74, 75]. The role of auxin during the development of the medial and lateral domains of the gynoecium is less clearly defined, however recent studies suggest that auxin homeostasis mechanisms are likely to be distinct in medial and lateral domains [23, 75, 76].

To better analyze auxin homeostatic mechanisms during medial domain development, we examined the expression of 127 genes with an annotated function in auxin homeostasis. Of these 127 genes, 80 were expressed in our dataset and 60 were differentially expressed at a FDR of < 0.01 in the YFP +/- comparison, without applying a fold enrichment filter (Additional file 11: Table S10). The expression levels of *TRYPTOPHAN AMINOTRANSFERASE OF ARABIDOPSIS 1 (TAA1)* and *YUC4*, two genes encoding proteins in the auxin synthetic pathway, were strongly enriched (> 4 fold) in the YFP positive samples as was predicted from previously published expression patterns indicating enriched expression within the medial portions of the gynoecium [33, 47, 77–80]. Within the *PINFORMED (PIN)* family of polar auxin transporters, the expression levels of *PIN1, PIN3* and *PIN7* were significantly enriched in the YFP-positive sample (Additional file 11: Table S10). This is consistent with the reported expression patterns at the protein level of these *PIN* transporters within the medial domain of the gynoecium [23, 76, 81, 82].

Auxin induces gene expression through a family of transcription factors called *AUXIN RESPONSE FACTORS (ARFs)* [74]. At a fold change level of 1.5 fold and FDR of <0.01, ten *ARFs* were enriched in the YFP-positive sample *(ARF1, ARF2, ARF3/ETTIN, ARF4, ARF5, ARF6, ARF7, ARF8, ARF16* and *ARF18)*, while no ARFs were identified as depleted in the YFP-positive sample (Additional file 11: Table S10). Our data suggests these *ARF* family members may be preferentially expressed in the medial domain and play a role during development of this meristematic tissue. Previous studies have documented gynoecial developmental defects in *arf3/ettin* mutants [4] as well as *arf6 arf8* [83, 84] double mutants. Interestingly, the levels of the precursor transcripts for two *TRANS-ACTING SIRNA3 (TAS3)* genes (At5g49615 and At3g17185) were significantly reduced (FDR <0.01) in the YFP-positive sample (Fig. 7b Additional file 9: Table S8). The trans-acting siRNAs that are encoded by the *TAS3* genes negatively regulate the levels of *ARF2, ARF3*, and *ARF4* transcripts [85]. Thus the enrichment of *ARF2 ARF3* and *ARF4* transcript levels in the *SHP2-expression* domain may in part be due to a reduction in the level of expression of the TAS3-encoded tasi-RNAs in the medial domain.

### *SQUAMOSA PROMOTER BINDING PROTEIN-LIKE3 (SPL3)* and the cis-NAT antisense gene At2g33815

The *SQUAMOSA PROMOTER BINDING PROTEIN-LIKE (SPL)* genes function in the regulation of the transition from juvenile to adult growth phases, and regulation of shoot regenerative capacity [86–89]. In our study, the expression of *SPL3* (At2g33810) was more than four fold lower in the YFP-positive sample relative to the YFP-negative sample. *SPL3* encodes a DNA-binding protein directly regulating *APETALA1* (At1g69120), a key regulator of floral-meristem-identity specification [90]. Interestingly, the expression of the cis-NAT antisense gene At2g33815, complementary to portions of the *SPL3* gene, was also significantly reduced ~4.5 fold in the YFP-positive sample (Cufflinks data in Additional file 4: Table S3 and Additional file 9 Table S8). This is perhaps in contrast to the expected pattern of expression, where the expression levels of the targeted *SPL3* transcript might be expected to go up as the levels of cis-NAT antisense At2g33815 go down. The expression of another regulator of *SPL3* activity, the *miRNA157D* (At1g48742), was also significantly reduced in the YFP-positive samples. The *miRNA157D* reduces translation of the *SPL3* transcript by acting through a *miRNA156/157-responsive* element in the *SPL3* 3’UTR [89, 91]. These data suggests that the *miRNA156/157/SPL* module may act during medial domain development and may be regulated by the cis-NAT antisense gene At2g33815. A complete list of differentially expressed natural-antisense, transposable-element and other non-protein coding transcripts identified as differentially expressed by Cufflinks, DESeq2 and edgeR is found in Additional file 9: Table S8.

### Protoplasting-induced stress genes

While the predominant focus of this work was to perform transcriptomic analysis in medial-domain-enriched cells (YFP+/-), transcripts induced by the protoplasting and sorting process (all-sorted/non-sorted) were also identified (Methods). To facilitate the visualization of all samples, we generated an interactive 6-way Venn diagram using the web-based tool ‘InteractiVenn’ [92]. By uploading the Additional file 12 to InteractiVenn [93], mousing over, and clicking on the numbers in the Venn diagram, researchers will find gene ID from DEGs between YFP+/- and all-sorted/non-sorted samples (3 programs and two comparisons). As expected, when comparing such different types of samples (all-sorted/non-sorted and YFP+/-), few DEGs (26) overlapped across the 6 samples (Fig. 8). The lack of overlap of DEGs across the entire experiment indicates that the YFP+/- DEGs reported here are not a result of protoplasting-stress-induced processes.

**Figure 8.**
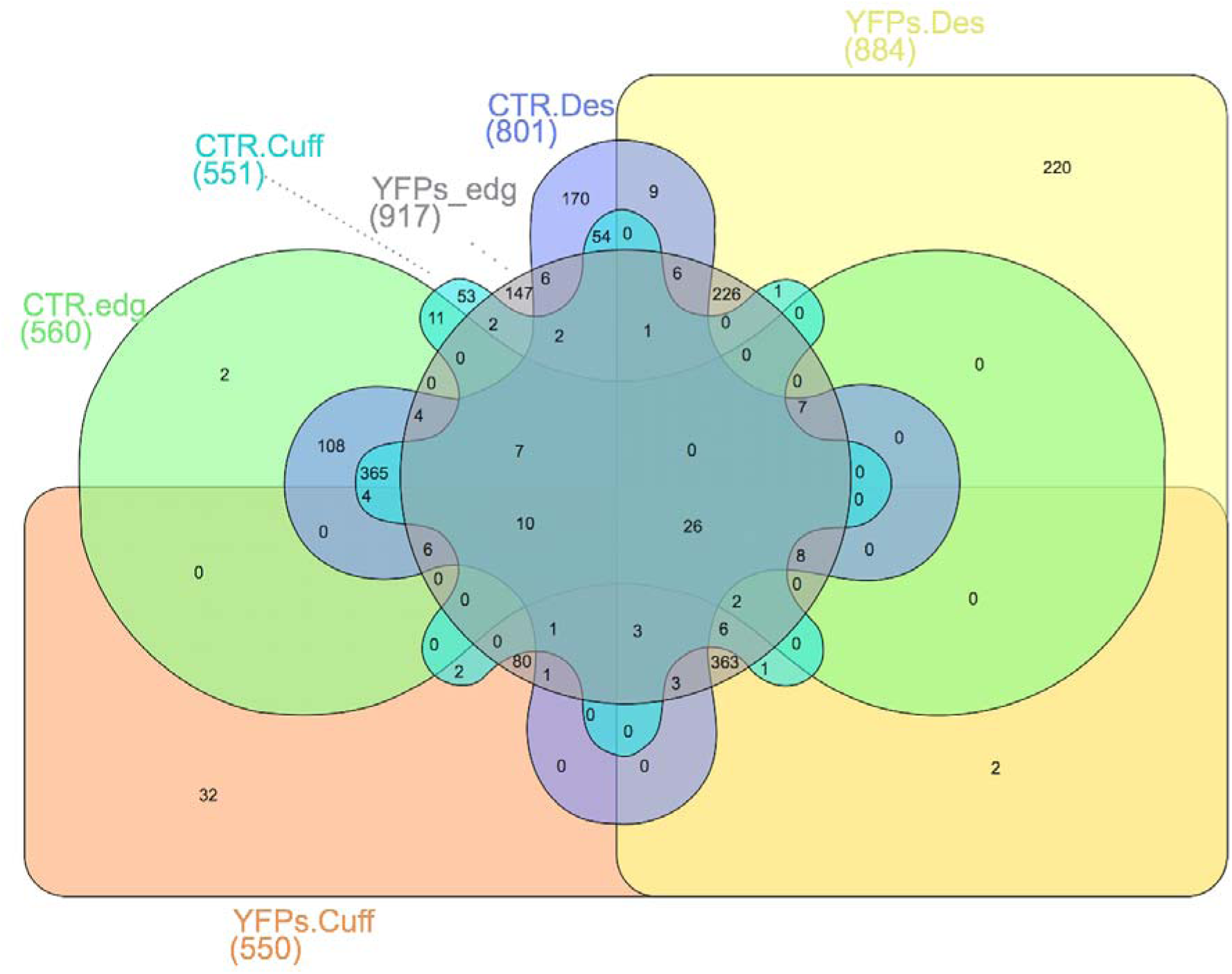
Six-way venn diagram image showing detailed overlap from all the differentially expressed gene (DEGs) datasets. The total number of DEGs under each condition and for each program are indicated in parentheses. CTR= DEGs between all-sorted/non-sorted and YFPs= DEGs between YFP+/-. Cuff= Cufflinks, edg=edgeR, Des=DESeq2. The interactive tool can be accessed online using the ‘InteractiVenn’ webtool [93] and uploading Additional file 12.

When comparing the protoplasting-induced gene set from this work (all-sorted/non-sorted DEGs) with those induced due to FACS-sorting methodology in shoot apical meristem by Yadav *et al*. [13] and in roots as reported by Birnbaum *et al*. [16] few DEGs were found in common (seven across all datasets) (Additional file 1: Figure S4) indicating that different tissues and/or different protoplasting techniques generate different sets of protoplast-induced gene-expression changes. Thus, appropriate controls should be included to control for condition-specific protoplasting-induced geneexpression changes.

## Conclusions

Despite the importance of the gynoecial medial domain in ovule development, no domain-specific transcriptome has been previously reported, mainly, due to the difficulty of isolating the meristematic cells from which ovules are derived. In this work, we developed a novel FACS-based system using the *SHP2*-expression-domain-specific using a GAL4/pUAS-based two-component system that, when combined with flower synchronization and flow cytometry, allowed for the efficient isolation of medial-domain cells expressing *SHP2*. The quality and quantity of biological samples that can be recovered with our system enables cell-type and strand-specific RNA-seq transcriptomic analysis and opens up possibilities for small RNAs, metabolomic and proteomic analyses [94–98]. This approach, coupled with high-throughput RNA-sequencing, has yielded a unique and novel snapshot of the gynoecial medial domain transcriptome and a set of candidate regulators of medial domain development for future functional analysis.

## Methods

### Construction of *pSHP2-GAL4//pUAS-3xYpet* dual construct lines

The *SHP2* promoter fragment was amplified from Columbia wild type genomic DNA using the primers proSHP2gwF1 (5’CACCATCTCCAACGCATTGTTACG3’) and proSHP2gwR1 (5’CATTTCTATAAGCCCTAGCTGAAG3’). This fragment contains the sequences from -2170 to +1 relative to the *SHP2* ATG and includes the 5’UTR, the first intron and the first Met codon of *SHP2*. This promoter previously was shown to mimic the endogenous *SHP2* expression pattern [22]. This genomic fragment was cloned into the pENTR/D-TOPO vector (Invitrogen) to create plasmid LJ001, and then shuttled via gateway LR reaction (Invitrogen) into the destination vector JMA859 (i.e. pEarleygate303-GAL4) to create plasmid AAS003. Transgenic Arabidopsis lines were created by *Agrobacterium*-mediated transformation of the AAS003 plasmid into the S. No. 1880 seed stock that contained the pGWB2-pUAS-3xYpet responder construct (see below) generating the pSHP2-GAL4; pUAS-3xYpet dual construct line (S. No. 1896), referred to as pSHP2-YFP. The pSHP2-YFP plants were crossed to the *ap1 cal1* Wellmer floral induction system [31] as described below.

JMA859 (pEarleygate303-GAL4) is a modified pEarleygate303 [99] plasmid in which the reporter was replaced by the coding sequences from the GAL4 yeast transcriptional activator. To achieve this, pEarleygate303 was cut with NcoI (New England Biolabs) and SpeI (New England Biolabs). Then fusion PCR was used to create the insert that fused the *GAL4* sequences to the deleted portions of pEarlygate303. This required three PCR reactions: 1st PCR with primers pEarl303NcoIFor

(5’TGGCCAATATGGACAACTTCT3’) and pEarl303Rev_GAL4 (tale) (5’ATGGAGGACAGGAGCTTCATACACAGATCTTCTTCAGAGA 3’); 2nd PCR with primers GAL4F_pEarl303 (tale)

(5 ’TCTCTGAAGAAGATCTGTGTATGAAGCTCCTGTCCTCCAT3 ’) and GAL4Rev_SpeI (5’ CCGGACTAGTCTACCCACCGTACTCGTCAA3’), and then a fusion PCR joining these two fragments using the external primers to amplify. The product of the fusion PCR was double-digested with NcoI/SpeI and ligated into NcoI/SpeI-cut pEarleygate303.

JMA382 (pUAS-pGWB2) was created from pGWB2 [100] by replacing the p35S sequences in pGWB2 with *pUAS* sequences (HindII/XbaI sites used). A Gateway LR reaction was then used to move the *3xYpet* cassette from JMA710 (pENTR/D-TOPO-3xYpet) into JMA382, creating vector JMA721 (i.e. pGWB2-pUAS-3xYpet). Homozygous single insertion-site transgenic lines harboring JMA721 were then generated (S.No 1880).

### Plant material

In a wild-type inflorescence, cells expressing *SHP2* represent a small percentage of the total cells. Additionally, wild-type inflorescence contains a full range of developmental series of floral stages. The Wellmer floral synchronization system [31] was used to maximize the amount of gynoecial tissue from floral stages 6-8 [12]. The Wellmer group kindly provided *pAP1-AP1::GR; ap1; cal* seeds (KanR in Ler background - S.No. 1927). *pSHP2-GAL4; pUAS-3xYpet* dual construct plants (S.No. 1896) were crossed to *pAP1-AP1::GR; ap1; cal*. Lines homozygous for *er, ap1, cal* and the transgenes were selected in F2 and F3 generations (generating S. No. 2060). Because of the mixed ecotype cross (Col and Ler), lines that were *erecta* homozygous mutant and gave consistent YFP expression pattern and consistent inducibility of the *AP1-GR* activity were selected before the generation of protoplasts. Plants were grown under constant light and temperature at 22 °C to minimize circadian transcriptional fluctuations. To induce flowering in the transgenic plants, 20 μm of the synthetic steroid hormone dexamethasone (DEX) (Sigma, USA) in 0.015 % silwet was applied directly (spray application) ~30 days after planting [31]. Inflorescences were collected for protoplast generation ~ 120 h after DEX-induced floral synchronization. When collecting samples for protoplast preparation, 5-6 inflorescence heads were fixed for chloral hydrate clearing and DIC microscopy to determine the developmental stages of the flowers of the inflorescence samples. Additionally, before protoplasting, whole inflorescences were also collected and frozen immediately in liquid nitrogen for analysis of the transcriptional starting state of the non-protoplasted tissue (non-sorted samples, see Experimental design).

### Experimental design

Material for RNA samples was gathered from batches of plants grown at one-week intervals to generate biological replicates (material from each week was considered as a biological replicate). To reduce variability between bioreplicates due to environmental heterogeneity within the growth chamber, each bioreplicate was drawn from a pool that contained plants grown within three different chamber positions. Four biological replicates of each of four tissue samples (YFP-positive, YFP-negative, all-sorted, and non-sorted) were collected (16 samples total). Whole inflorescences were collected for non-sorted samples and immediately frozen in liquid nitrogen before RNA isolation (i.e. these samples were not subjected to protoplasting nor FACS-sorting). The all-sorted samples represented the total population of protoplasts that come off the FACS machine after debris and broken cells are removed based on sorting gates (Additional file 1: Figure S1). The YFP-positive and YFP-negative protoplast populations are processed equivalently to the all-sorted samples except that a final FACS-sorting gate is used to divide the all-sorted protoplasts into YFP-positive and YFP-negative samples (Additional file 1: Figure S1). RNA was isolated from these three protoplast populations, as well as from entire non-protoplasted inflorescences (“non-sorted”). The YFP-positive, YFP-negative and all-sorted samples were prepared and collected as described below (Protoplast recovery and cell sorting).

### Protoplast recovery and cell sorting

Protoplasts from the S. No. 2060 plants were generated according to the protocol of Birnbaum *et al*. [16], with adaptations for inflorescence plant material. Inflorescences (~200) were hand-collected with forceps and/or scissors and chopped with a “Personna double edge prep blade” (American Safety razor company; 74-002) within a 15 min period. Cell-wall polysaccharides were digested by immersing the chopped plant material in 10 ml of filter-sterilized solution B in a 50 ml falcon tube. Solution B (prepared according to Birnbaum *et al.)* is prepared from Solution A (10 mM KCl, 2 mM MgCl_2_, 0.2M MES, 600 mM Mannitol) to which cell wall digesting enzymes were added [final concentrations of 1.5% Cellulase (Yakult, Japan), 1% Pectolyase (Yakult, Japan) and 1% Hemicellulase (Sigma, USA)]. This mixture is then dissolved by gently swirling, covered in foil, and warmed in a water bath at 55 °C for ten minutes to inactivate DNAses and proteases. After cooling to room temperature, CaCl_2_ (2 mM final) and BSA (0.1% final) were added and the solution was filter-sterilized through a 25-micron filter.

After 1 h of incubation at room temperature with occasional gentle agitation, 10 ml of the protoplast-rich solution B was filtered through a 70-micron filter basket to a 50 ml falcon tube. A 10 ml rinse of solution A was applied directly to the material left in the 70-micron filter basket to rinse through any protoplasts left behind. Protoplasts were spun at 500 g, 10 °C for 10 min; the majority of the supernatant was removed by aspiration being careful not to disturb the protoplast pellet which is typically not tightly compacted. Protoplasts were resuspended in 25 ml of Solution A as a rinse step to remove cell-wall-digesting enzymes. Protoplasts were filtered again through a 50-micron filter mesh to a new tube adding 8 ml of solution A to again rinse through any protoplasts stuck in the filter. Protoplasts were then spun again at 500 g for 10 min. The majority of the supernatant was removed leaving 2 ml of the protoplasts in solution after the second centrifugation step. Propidium Iodide (5 micrograms/ml final) was added to the protoplasts (to allow separation of broken protoplasts) and a final filtering step though a 30-micron mesh filter (CellTrics, Partec) was carried out before loading onto the FACS machine.

Flow cytometry through FACS-sorting (Moflo XDP; Beckman Coulter Inc.) was used to isolate the YFP expressing cells from the total pool of cells. The FACS machine was equipped with a cooling device (set to 10 °C) and fitted with a 100-μm nozzle. Protoplasts were sorted at a rate of up to 10,000 events per second at a fluid pressure of 25 psi. Four sorting gates were set in an effort to collect the cleanest set of protoplasts and to eliminate debris and broken cells. A first gate based on size and granularity using side-scatter (SS) and forward-scatter (FS) parameters was used to select for intact protoplasts. Then a second gate was used to select for single cells and remove “doublets”. A third gate was used to select for cells that were negative for propidium iodide (PI) signal, as broken protoplasts and debris are preferentially stained by PI, which is excited by the 488 nm laser and emits at 617 nm. The total population of protoplasts that came off the FACS sorter machine after these gates constituted the all-sorted sample. In parallel, the YFP-positive protoplasts and YFP-negative protoplasts were separated into two collection tubes using the gates described above and one additional sorting gate based on the level of emission intensity in the green channel (529nm/28nm filter). Preliminary experiments with protoplasts that did not express the YFP transgene were used to set this gate and determine the levels of auto-fluorescence of the protoplasts. Protoplasts were collected directly into 14 ml tubes containing 4 ml of Trizol (Invitrogen/Life Technologies) and occasionally agitated during the approximately 40 min of sorting required to collect the protoplasts. Trizol was the method of choice as it maintains a high level of RNA integrity during tissue homogenization while also disrupting and breaking down cells and cell components. In order to minimize artifactual changes to transcript levels, the entire process of cell wall digestion, protoplast generation and FACS-sorting was kept under three hours. This procedure typically yielded between 300,000 and 500,000 YFP-positive protoplasts. These YFP-positive protoplasts typically represented approximately 0.5% of the total FACS sorting events. On average from four sorting trials representing four biological replicates, the number of cells collected and processed for each sample was: 575K for the YFP-positive, 1000K for the YFP negative and 493K for all-sorted samples.

### RNA extraction and quantitative RT-PCR

Total RNA was extracted from sorted protoplasts collected in Trizol (keeping a 3:1 ratio of Trizol to sorted cells) and by modifying the Plant RNeasy Mini Kit, Qiagen protocol, as follow: collected cells in Trizol (4 ml total) were vortexed for 5 min at room temperature (RT) and 1 ml of chloroform (Sigma) was then added. The solution was vortexed again for 1 min at RT and centrifuged at 4,000 rpm for 10 min at 4 °C to separate phases; RNA from the aqueous phase (top layer) was carefully sucked up and mixed with 700 μl of Qiagen RLT buffer (Plant RNeasy Mini Kit, Qiagen) and 7μl of B-Mercaptoethanol (Sigma). 500 μl of 100% ethanol was added, solution was then transferred to a Qiagen MinElute column (Plant RNeasy Mini Kit, Qiagen) and spun in a 2 ml microfuge tube for 15 sec at ~ 10,000 rpm. 500 μl of RPE (Plant RNeasy Mini Kit, Qiagen) was added to the spin column, spun for 15 sec at ~10,000. 750 μl of 80% ethanol was added to the MinElute column and spun at ~10,000 rpm for 15 sec (twice) to ensure removal of all guanidine salts that may inhibit downstream applications. A final 5 min spin at top speed with the cap off was performed to remove trace amounts of ethanol. Total RNA was then eluted with 10 μl of RNAse-free water. A second elution was performed with another 10 ul of RNAse-free water. It is worth noting that one biological replicate (4^th^ biological replicate) from the YFP-positive protoplasts was lost at this point, leaving only 3 biological replicates for this tissue sample and yielding a total number of 15 samples sequenced in two lanes and used for the experiment.

Prior to high-throughput sequencing, quantitative RT-PCR (qRT-PCR) was conducted on YFP-positive and YFP-negative samples using the 2^−ΔΔCT^ method as suggested by Schmittgen and Livak [101] to assess relative gene expression of specific medial domain markers, *SHATTERPROOF2* and *NGATHA1*. Total isolated RNA was quantified using fluorometric quantitation (Qubit RNA Assay Kit, Life Technologies, Inc.) for both YFP-positive and YFP-negative samples [~ 100 ng]. SuperScript III First-Strand Synthesis System (Invitrogen/Life Technologies) was used to generate cDNA (cDNA diluted 1:4 prior qRT-PCR analysis) from total RNA. qRT-PCR experiment assay was performed (Thermal Cyclers from Applied Biosystems) using a SYBR green mix (QuantiTect SYBR Green PCR Kits, Qiagen). Three biological replicates of the YFP-positive and YFP-negative samples were included and each biological replicate was assayed in triplicate. The expression levels of the *ADENINE PHOSPHORIBOSYL TRANSFERASE1 (APT1)* (At1g27450) gene was used for normalization.

### Barplots

Barplots graphs were constructed using the ‘R’ package bear [102] and plyr [103] to calculate mean, standard error and confidence intervals and ggplot2 [104] to generate the plots.

### Library preparation and mRNA sequencing

Total RNA isolated was quantified using fluorometric quantitation (Qubit RNA Assay Kit, Life Technologies, Inc.) and RNA quality was assessed using Agilent 2100 Bioanalyzer (Agilent). The RNA integrity number (RIN) for the 15 samples was higher than 7.3, which is above the Illumina threshold for library construction (> RIN 7). Strand-specific cDNA libraries were constructed from approximately 100 ng of total RNA using a NEB Ultra Directional Library Prep Kit for Illumina (New England Biolabs). The average size of the cDNA fragments was ~ 250 bp. The 15 bar-coded libraries were pooled and single-end sequencing was performed in a HiSeq 2500 Illumina (Illumina, Inc.) with ‘HiSeq SR Cluster Kit v4’ for the flow-cell and ‘HiSeq SBS v4’ for sequencing reagents. cDNA libraries were sequenced in 125-cycle plus 7-cycle for multiplexed samples. Sequencing was performed in two lanes of a flow-cell; all 15 libraries were sequenced twice and the results from the two independent lanes were analyzed as technical replicates. As no lane-specific effects were observed during data analysis, the reads from each lane were pooled for analysis of DEGs (see Table counts and technical replicates).

### Bioinformatics analysis

All bioinformatics analyses were performed on a server cluster with 128 GB (gigabytes) of RAM, 16 cores (CPUs) and Ubuntu Linux-Distribution 12.04 operating system using ‘Simple Linux Utility Resource Management’ (SLURM) queue management system at the Bioinformatics Research Center (BRC) at the North Carolina State University, Raleigh, NC, USA.

### Read Processing

Quality control and preprocessing of metagenomic data was performed using FastQC software [105]. Adapters and low quality sequences were filtered out with Ea-Utils software [106]. Reads with phred-like quality score (Q-score) > 30 and read length > 50-bp were kept and aligned against the TAIR10 Arabidopsis reference genome.

### Sequence alignment to the Arabidopsis genome

Splice junction mapper TopHat2 (version 2.0.10) [107] was used to align filtered RNA-seq reads to the *Arabidopsis thaliana* TAIR10 genome (Ensembl annotation) downloaded from the iGenome database [108]. Default parameters for TopHat2 were used except for strand specificity (--library-type=fr-firststrand) to match to the first strand of cDNA synthesized (anti-sense to the mRNA) and maximal intron length (--I 2000), as it has been shown that the large majority of the known introns are smaller than the selected threshold [96]. To align reads solely and exclusively against TAIR10 annotated gene models, the arguments ‘--T’ (transcriptome only) and ‘--no-novel-juncs’ (no novel junction) were also included. Uniquely mapped reads were extracted from the TopHat2 output binary (BAM) file using samtools [109] and selecting for the “NH:i:1” two-character string-tag. Only uniquely mapped reads were used for downstream analysis.

### Table counts and technical replicates

The ‘HTSeq: Analyzing high-throughput sequencing data with Python’ software [110] was used with default parameters except for the ‘stranded=reverse’ mode to generate tables-counts for downstream differential expression analysis for the ‘R’ packages edgeR [36] and DESeq2 [37].

Using edgeR, we assessed the gene level variance versus log gene expression level among technical replicates (corresponding to two lanes in the flow-cell of the Illumina HiSeq 2500). A linear-dependent Poisson distribution was observed for technical replicates (Additional file 1: Figure S5), in accordance with several studies [36], [67], [111]. Thus, differential gene expression analysis was performed using pooled technical replicates.

### Gene expression and differential gene expression

Gene expression and differential gene expression analysis was carried out using ‘R’ packages edgeR [36] and DESeq2 [37] and the Linux-based Cufflinks program (v2.2.1) (G option) [35], for differentially expressed genes and transcripts [35]. To facilitate future use of these datasets, all the expressed genes identified and their expression values (F/RPKM) in YFP+/- (Additional file 4: Table S3) and all-sorted/non-sorted (Additional file 5: Table S4) are included as supplementary material.

Filters were applied to determine if a gene was detected, abiding by the suggestions of statisticians and bioinformaticians [112], [113], [114], [115], [37], [116] as a means to enrich for true DEGs, to reduce type I error and to improve *P*-value adjustment. The edgeR function [36] ‘cpm’ (counts per million) was used to discard those genes whose cpm was lower than a threshold of 2 reads per gene in at least 3 biological replicates, as suggested in the edgeR vignette. For cufflinks, a minimum RPKM of 5 was set for a gene to be expressed, following Suzuki *et al*. criteria [117]. According to Sims *et al*. 80% of genes can be accurately quantified with FPKM > 10 [69]. DESeq2 performs independent filtering using the ‘results’ function, as described in the DESeq2 vignette [37]. An FDR cutoff of < 0.01 was used to determine differentially expressed genes in all three programs. The Gene Regulatory Information Server (AGRIS) was used to identify transcription families in the dataset [118]. Enriched categories of transcription factors within the set of “cleaned” 363 YFP+/- DEGs was assessed with the online Transcription Factor Enrichment Calculator tool [51].

### Venn Diagrams and heatmaps

Venn diagrams were constructed using the ‘R’ package VennDiagram [119] and the web-based tool package InteractiVenn [92]. Heatmaps were produced using the ‘R’ package pheatmap [120]. RPKM normalization by gene length and library size values were produced using the ‘rpkm’ function from edgeR [36]. To calculate gene length, a TAIR10 gene length list (CDS plus UTRs) was constructed by extracting length information from the TAIR10 GFF file with homemade Perl script. Genes with multiple isoforms were collapsed and length was calculated using the longest one. RPKM values were then calculated for clustering purposes and to have an intermediate point of comparison between Cufflinks, edgeR and DESeq2. Samples were clustered (default clustering) with parameters provided in the software. The ‘R’ package colorRamp [121] was used to produce a gradient of color values corresponding to gene-fold change values.

### Gene set enrichment analysis

Gene Ontology (GO) enrichment tests were performed using the ‘R’ package topGO [122], with the ‘classic’ algorithm (where each GO category is tested independently) and the ‘fisher’ statistic test for ‘biological processes’, ‘molecular function’ and ‘cellular component’. Enrichment analysis was performed separately for all the genes that were differentially expressed between the YFP+/- samples and between the all-sorted/non-sorted samples. Network analysis of GO terms was performed using BiNGO [123] plugin for Cytoscape [124]. GO terms for the 268 genes identified as depleted in the YFP-positive sample, as well cellular component (CC) and molecular function (MF) for the YFP+/-sample can be found in Additional file 7: Table S6.

### Dendrograms

The ‘R’ Dist function was used to compute a distance matrix using the spearman method (Spearman test rank correlation) and the ‘R’ Cor function to compute the variance of the matrix. To perform hierarchical clustering, the hclust function in ‘R’ was used. All statistical analyses were performed in ‘R’ v.3.0.2. Dendrogram plots were built using the ‘R’ ape package with edge.color = "blue".

### Confocal microscopy

Confocal microscopy was performed using a Zeiss LSM 710 (Carl Zeiss, Inc. Thornwood, NY), microscope model (Zeiss Axio Observer Z.1), objective type Plan-Apochromat 20x/0.8 M27. Z-stack intervals were set to 2 μm and the total thickness of the stack was 62 μm.

### Chloral hydrate clearing and Differential Contrast (DIC) Microscopy

Inflorescence samples were fixed in a solution of 9 parts ethanol: 1 part acetic acid for two hours at room temperature, and then washed twice in 90% ethanol for 30 min each wash. Inflorescences were transferred to Hoyer’s solution (70% Chloral hydrate w/v, 4% glycerol, 5 % gum Arabic) and allowed to clear for several hours to overnight. Samples were then dissected in Hoyer’s solution. The dissected inflorescence heads were mounted in Hoyer’s under coverslips and examined with DIC optics on a Zeiss Axioskop 2 to determine the floral stages.

### *In situ* hybridization

For *in situ* hybridization analysis, *Arabidopsis thaliana* Col-0 flowers were fixed and embedded in paraffin as described previously [11, 70]. Sections of plant tissue were probed with digoxigenin-labeled antisense and sense RNA probes (Roche). Probes corresponded to nucleotides +686 to +920 of *REM13* relative to the transcriptional start site of the CDS using the following oligos to amplify the template: REM13_ISH_Fwd 5’ AAAATAGAACGCGCATACCG 3’ and REM13_ISH_Rev 5’ TCGTGAACCAAACCGTGATA 3’. Hybridization and immunological detection were performed as described previously [11, 70].

### Data availability

Illumina sequencing raw data (fastq) have been submitted to the Gene Expression Omnibus (GEO) database (accession GSE74458).

**List of abbreviations**
bp: base pair
CMM: carpel margin meristem
DEG: differentially expressed gene
DEX: dexamethasone
FACS: fluorescence-activated cell sorting
FDR: false discovery rate
FPKM: fragments per kilobase of transcript per million mapped reads
GSEA: gene set enrichment analysis
LCM: laser capture microdissection
nt: nucleotide
PI: propidium iodide
qRT-PCR: quantitative RT-PCR
RPKM: reads per kilobase of transcript per million mapped reads
ssRNA-seq: strand-specific RNA sequencing
TMM: trimmed mean of M-values
TSS: transactional start site
UTR: untranslated region
YFP: yellow fluorescent protein

## Authors’ contributions

RGF, GHV, ES and SH coordinated and conceived of the study. AS, JV, LR built the GAL4/pUAS-based two-component reporter system. GHV, MFV and BS performed FACS-sorting, wet-lab and microscopy analyses. SM and LC performed *in situ* hybridization. GHV, QH and SH performed RNA-seq and related bioinformatic and computational analysis. GHV and RGF interpreted the data and drafted the manuscript. All of the authors revised and approved the final manuscript.

## Description of additional data files

The following additional data are available with the online version of this paper. Additional data files Figures S1-S5 contain sorting gates used to select YFP samples and the re-sorting of the YFP-positive cells to assess sample purity (Fig. S1), qRT-PCR enrichment of medial domain genes *SHP2* and *NGA1* and the gene *TUB* (FIG. S2), expression profiles for the 363 differentially expressed genes (FC >4, FDR <0.001) across all 4 samples (YFP-positive, YFP-negative, all-sorted, non-sorted) (Fig. S3), Venn Diagram comparison of stressed induced genes due to protoplast/FACS-sorting procedure (Fig. S4) and gene level variance versus log gene expression level among technical replicates (Fig. S5). Additional file 2: Table S1 containing summary RNA-seq data (number of reads, mapped reads, uniquely mapped, etc.). Additional file 3 Table: S2 contains differentially expressed genes (DEGs) from the YFP+/- and all-sorted/non-sorted comparison. Additional file 4: Table S3 and Additional file 5: Table S4 contain all the expressed genes identified with three different programs between all the YFP+/- samples and all-sorted/non-sorted samples, respectively. Additional file B: Table S5 corresponds to raw high-throughput count data for YFP+/- and all-sorted/non-sorted comparison. Additional file 7: Table S6 contains the set enrichment analyses (GSEA) for YFP+/- and all-sorted/non-sorted comparison, including Biological Process (BP), Molecular Function (MF) and Cellular Component (CC). Additional file 8: Table S7 lists the transcription factors families identified in the DEGs from YFP+/- and their statistical enrichment Additional file 9: Table S8 contains isoforms expression, regulation of gene expression by alternative promoters and antisense transcripts identified by Cufflinks, edgeR and DESeq2. Additional file 10: Table S9 corresponds to the expression profile (RPKM) of the 86 genes described by Reyes-Olalde *et al*. [8] expressed in the medial domain. Additional file 11: Table 10 contains hormone (Auxin, GA, Ethylene) related-genes present in our dataset. Additional file 12 is the datafile to upload the web-based tool package “InteractiVenn”.

## Acknowledgements

We thank Sarah Schuett (CVM, flow cytometry facility, NCSU) and the Genomic Sciences Laboratory (GSL) research facility (NCSU) for library preparation and Illumina sequencing. We also would like to thank William Thomson (NCSU) and Emily Wear (NSCU) for FACS-sorting assistance; Frank Wellmer and Diarmuid O’Maoileidigh (Smurfit Institute, Trinity College of Dublin, Ireland) for the *pAP1:AP1:GR ap1 cal1* floral synchronization system; Colleen Doherty (NCSU) for help with Cytoscape/BiNGO analysis; Maria Angels De Luis Balaguer (NCSU) for assistance with cross-platform expression correlation approaches; José Alonso and Ross Sozzani (NCSU) for thoughtful comments on the manuscript; Jigar Desai, Dmitry Grinevich and Colleen Doherty (NCSU) for help with the transcription factor enrichment analysis. Aureliano Bombarely (Virginia Tech) for help with GO terms analysis; and Eva Johannes (CMIF, Molecular Imaging Facility, NCSU) for laser scanning confocal microscope assistance. This work was funded by a grant from the National Science Foundation to RGF and SH (NSF IOS-1355019) and the FP7-PEOPLE-2013-IRSE FRUIT LOOK program (to ES, LC and RGF).

## Supplementary Figure

**Additional file 1: Figure S1.**
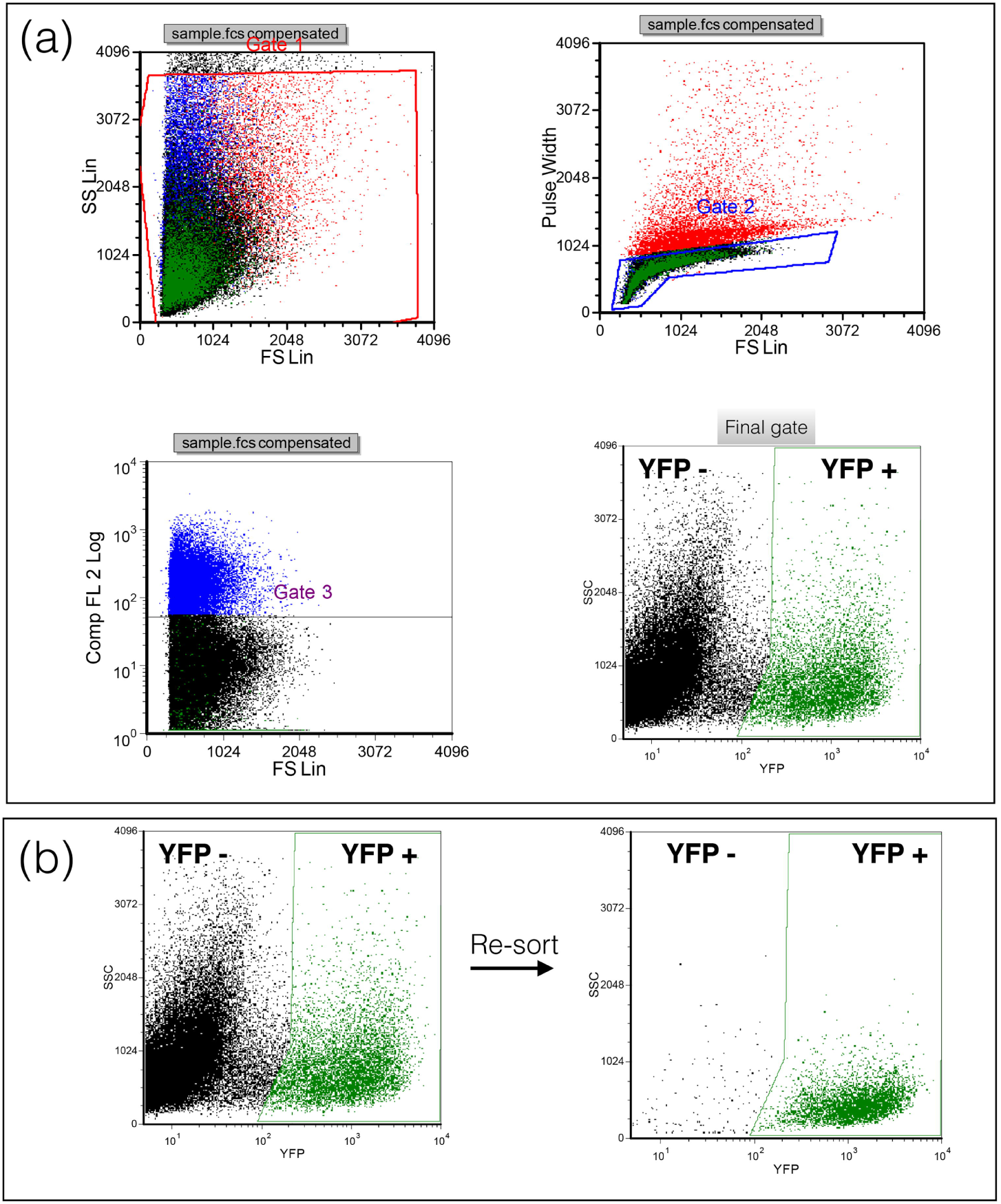
Sorting/re-sorting YFP-positive cells yielded high cell-type specific purity. Panel **a** Sorting gates used to select for fluorescent YFP-positive, YFP-negative and all-sorted protoplasts. **b** Re-sorting YFP-positive protoplasts to assess sample purity.

**Additional file 1: Figure S2.**
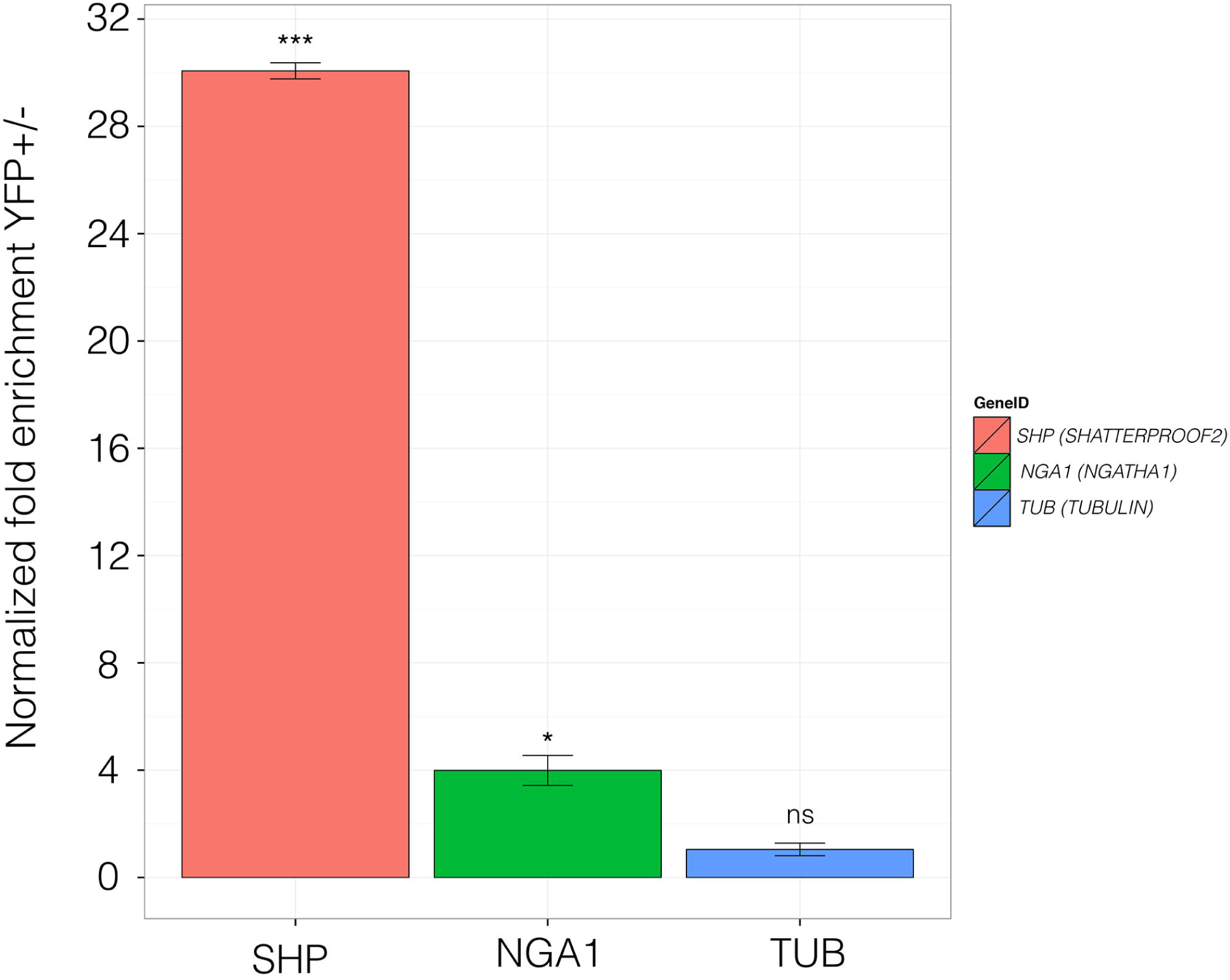
Enrichment of medial-domain expressed genes, *SHATTERPROOF2 (SHP2)* and *NGATHA1 (NGA1)*, assessed by quantitative real time PCR (qRT-PCR) from FACS-sorted protoplasts corresponding to YFP-positive and YFP-negative samples. The differences in the level of expression between the YFP positive and YFP negative samples transcripts were statistically significant for *SHP2* and *NGA1, p* <0.001 and *p* <0.05, respectively. The difference in the levels of expression for the *TUBLIN6* gene was not found to be statistically significant (*p* = 0.4). The expression levels of the *ADENINE PHOSPHORIBOSYL TRANSFERASE1* (APT1) transcript was used for normalization.

**Additional file 1: Figure S3.**
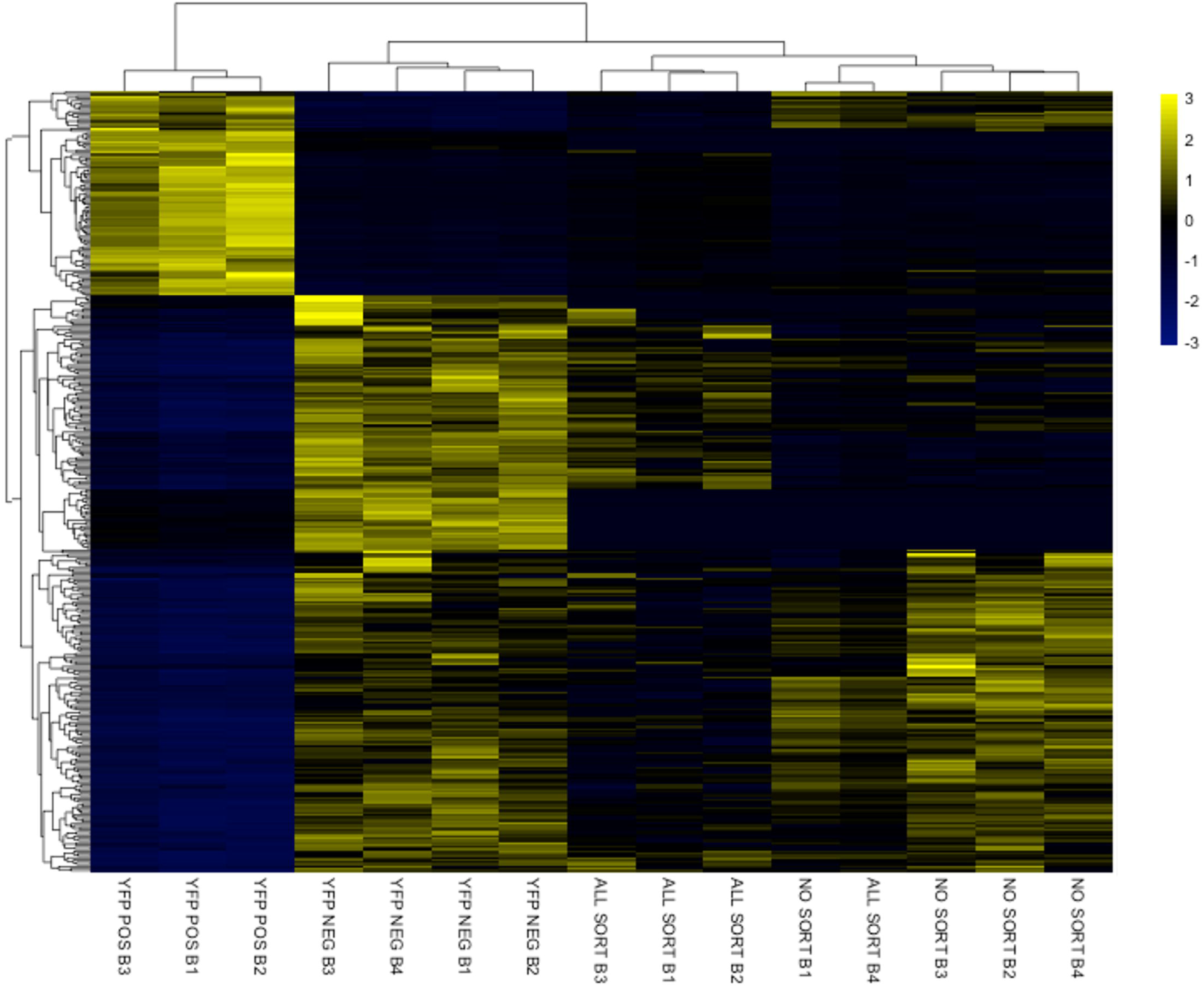
Transcriptional profile in Reads Per Kilobase of transcript per Million mapped reads (RPKM) of the 363 selected differentially expressed genes across all the samples used in this study (YFP POS = YFP-positive, YFP NEG = YFP-negative, ALL SORT = all-sorted, and NO SORT = non-sorted). Biological replicates are indicated as B1, B2, B3, and B4.

**Additional file 1: Figure S4.**
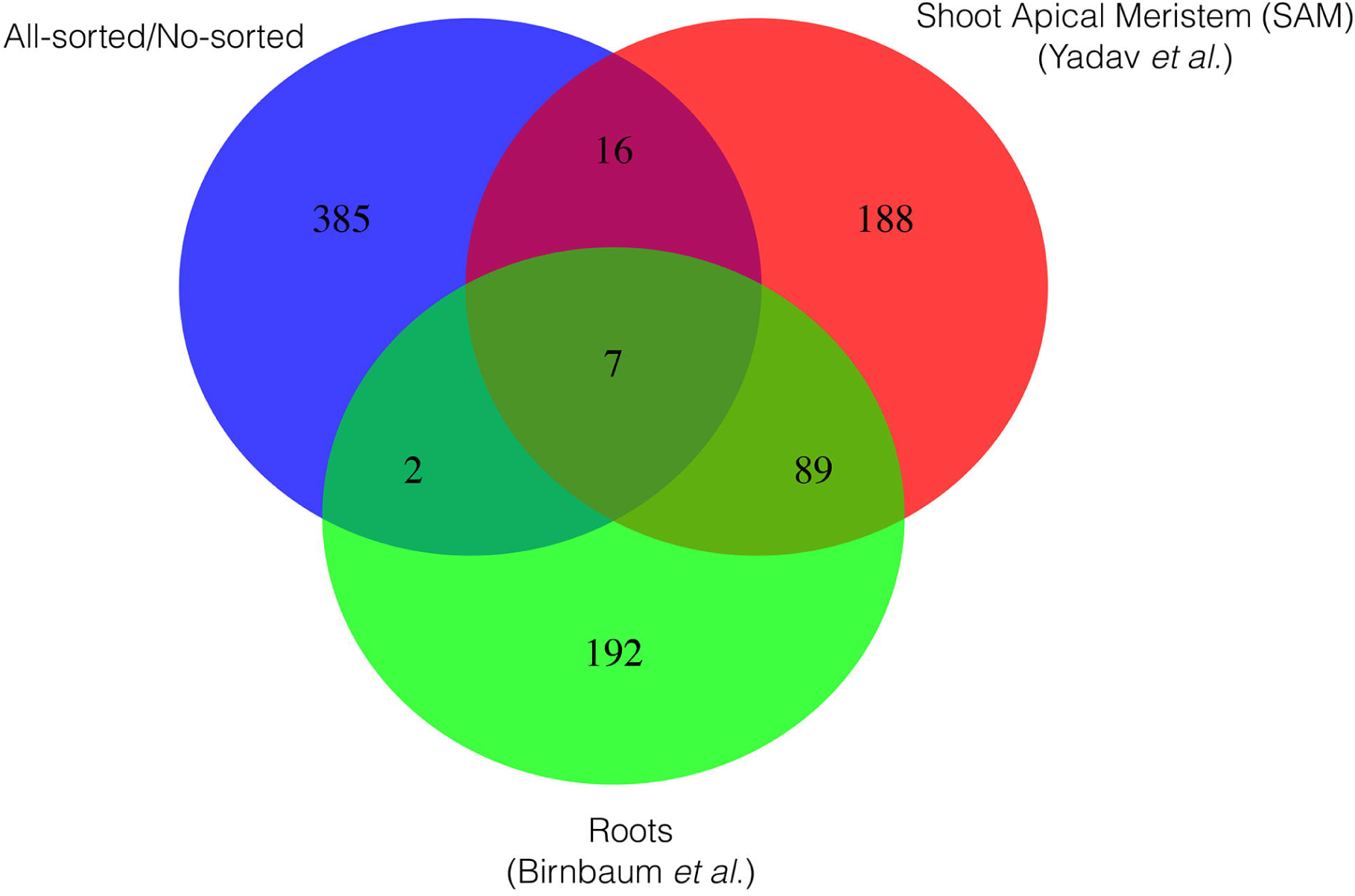
Venn Diagram of stressed induced genes due to the protoplast/FACS-sorting procedure including Shoot Apical Meristem (SAM) samples [13] and root samples [16].

**Additional file 1: Figure S5.**
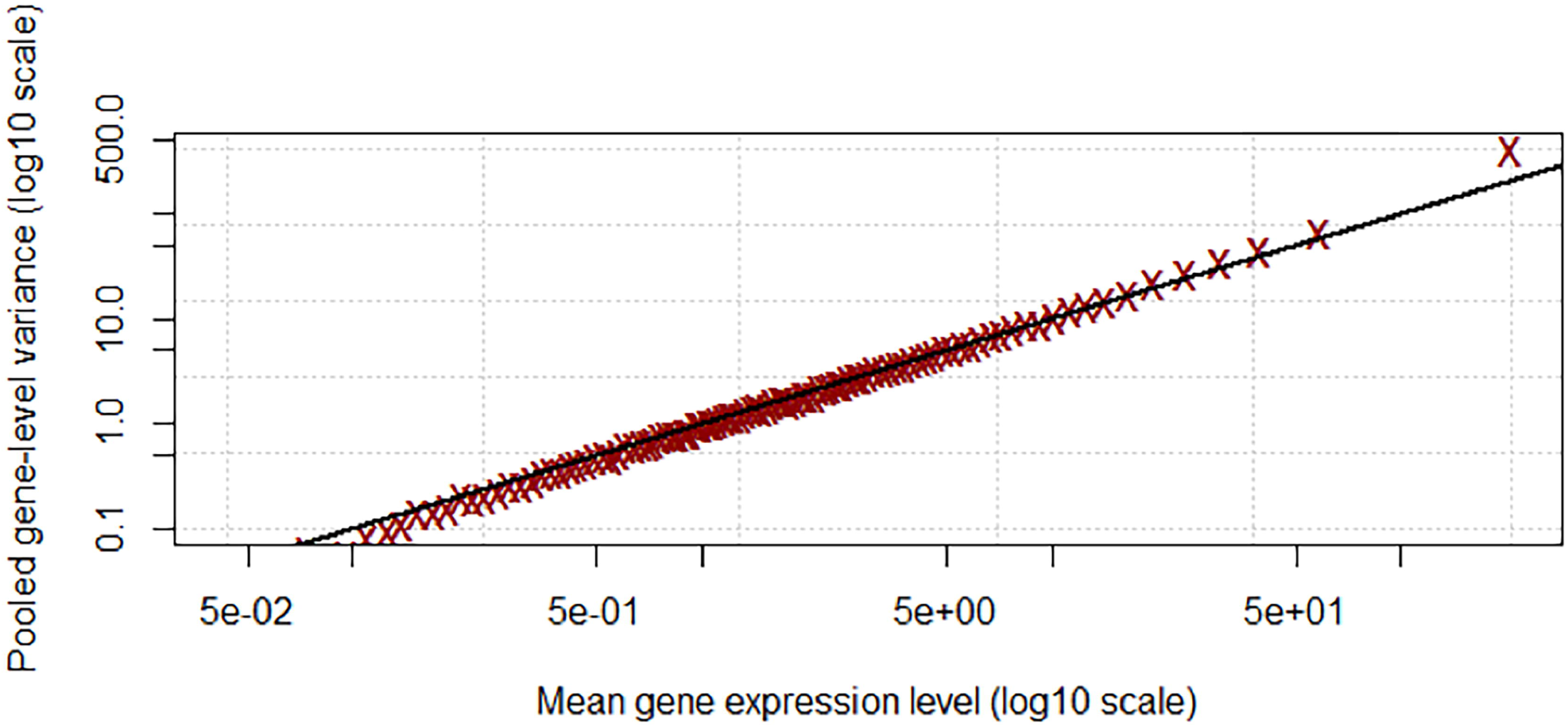
Plot of pooled gene level variance versus log gene expression level among technical replicates.

## Supplementary Tables

**Additional file 2: Table S1.** Sample names are listed in the first column (YFP-positive, YFP-negative, all-sorted, non-sorted). The 2^nd^ and 3^rd^ column correspond to million DNA reads (MR) in each technical (T1 and T2) and biological replicate (B1, B2, B3, B4), respectively. Filtered reads (Q30L50) per sample are found in the 4^th^ column and 5^th^ column (technical and biological replicate, respectively). The 6^th^ column indicates the number of the MR filtered out after removing adapters and low quality sequences. The 7^th^ and 8^th^ columns indicate MR mapped to the Arabidopsis TAIR10 genome. The 9^th^ and 10^th^ column include uniquely mapped reads to the Arabidopsis TAIR10 genome and 11^th^ the MR that aligned more than in one place in TAIR10 genome. The 12^th^ and 13^th^ column represent reads mapped to only annotated TAIR10 genes (technical and biological replicates). The 14^th^ and 15^th^ columns correspond to uniquely mapped reads to annotated genes.

**Additional file 3: Table S2.** Table 1 - List of 363 differentially expressed genes (YFP+/-) with FDR <0.001 and fold change > 4 co-identified by three independent software packages (Cufflinks, DESeq2, and edgeR). Table 2 - List of 410 differentially expressed genes (all-sorted/non-sorted) with FDR <0.01 and fold change > 4 co-identified by Cufflinks, DESeq2 and edgeR. Table 3 - List of 48 genes differentially expressed in both the YFP+/- comparison and the non-sorted/all-sorted comparison.

**Additional file 4: Table S3.** All expressed genes, FDR, p-values, and their expression levels (in Reads Per Kilobase of transcript per Million mapped reads (RPKM)) identified with Cufflinks (Table 1), DESeq2 (Table 2), and edgeR (Table 3) between YFP+/YFP-samples.

**Additional file 5: Table S4.** All expressed genes, FDR, p-values, and their expression levels (in Reads Per Kilobase of transcript per Million mapped reads (RPKM)) identified with Cufflinks (Table 1), DESeq2 (Table 2), and edgeR (Table 3) between all-sorted/non-sorted samples.

**Additional file 6: Table S5.** Table counts produced with ‘HTSeq: Analyzing high-throughput sequencing data with Python’ software (HtSeq) for all the samples (YFP-positive, YFP-negative, all-sorted, and non-sorted) including all biological replicates. Table 1 (YFPs) - YFP-positive (YFP_POS) and YFP-negative (YFP_NEG), Table 2 - allsorted (ALL_SORT) and non-sorted (NO_SORT). Biological replicates are indicated as B1, B2, B3, B4.

**Additional file 7: Table S6.** List of GO terms including biological process (BP), cellular component (CC) and molecular function (MF) categories identified in the gene set enrichment analyses (GSEA) for the enriched differentially expressed genes (95) and for the depleted DEGs (238) between YFP+/- from the 363 DEGs list (\*\**p*-value <0.01). DEGs. Over-represented GO categories (\*\**p* < 0.01) between all-sorted/non-sorted are also presented in Table 2, labeled as “ALL vs. NON_SORTED”.

**Additional file 8: Table S7.** Table 1 - List of 75 transcription factors and associated transcription factor family designations identified in the 363 YFP+/- DEGs. Table 2 - enriched categories of transcription factors within the set of the 363 DEGs using the online Transcription Factor Enrichment Calculator tool [51].

**Additional file 9: Table S8.** Table 1-Differential isoform expression that are significant (a=0.01) between the YFP+/- comparison and non-significant between the all-sorted/non-sorted comparison. Table 2- List of genes differentially expressed (a=0.01) by TSS/alternative promoter usage. Differentially expressed non-coding RNAs between the YFP+/- samples are reported in Table 3, 4, 5 for edgeR, DESeq2 and Cufflinks, respectively.

**Additional file 10: Table S9.** Expression profiles in Reads Per Kilobase of transcript per Million mapped reads (RPKM) of the 86 genes described by Reyes-Olalde *et al*. [8] in our dataset; YFP-positive (YFP_POS), YFP-negative (YFP_NEG), non-sorted (NO_SORT), all-sorted (ALL_SORT) and expressed in the medial domain of the arabidopsis gynoecium. Data from Mantegazza *et al*. [54] are also included; inflorescence meristem (IM_AVG), floral meristem (FM_AVG) and floral stage 3 (ST3_AVG).

**Additional file 11: Table S10.** List of genes associated with auxin, gibberellin and ethylene, their expression values, fold changes, and significance in our dataset.

**Additional file 12:** Data file to upload the web-based tool package “InteractiVenn” [93].

## References

1. Seymour GB, Østergaard L, Chapman NH, Knapp S, Martin C: Fruit development and ripening. Annu Rev Plant Biol 2013, 64: 219–241.

2. Oram RN, Brock RD: Prospects for improving plant protein yield and quality by breeding. Australian Inst Agr Sci J1972.

3. Singh MB, Bhalla PL: Control of male germ-cell development in flowering plants. Bioessays 2007, 29: 1124–1132.

4. Sessions RA, Zambryski PC: Arabidopsis gynoecium structure in the wild and in ettin mutants. Development 1995, 121: 1519–1532.

5. Bowman JL, Baum SF, Eshed Y, Putterill J, Alvarez J: 4 Molecular Genetics of Gynoecium Development in Arabidopsis. Curr Top Dev Biol 1999, 45: 155–205.

6. Crawford BCW, Yanofsky MF: The Formation and Function of the Female Reproductive Tract in Flowering Plants. Curr Biol 2008, 18:R972-R978.

7. Sessions A: Piecing together the Arabidopsis gynoecium. Trends Plant Sci 1999, 4:296–297.

8. Reyes-Olalde JI, Zuñiga-Mayo VM, Chávez Montes RA, Marsch-Martínez N, de Folter S: Inside the gynoecium: at the carpel margin. Trends Plant Sci 2013, 18: 644–655.

9. Alvarez J, Smyth DR: CRABS CLAW and SPATULA Genes Regulate Growth and Pattern Formation during Gynoecium Development in Arabidopsis thaliana. Int J Plant Sci 2002, 163: 17–41.

10. Aichinger E, Kornet N, Friedrich T, Laux T: Plant stem cell niches. Annu Rev Plant Biol 2012, 63: 615–636.

11. Wynn AN, Rueschhoff EE, Franks RG: Transcriptomic Characterization of a Synergistic Genetic Interaction during Carpel Margin Meristem Development in Arabidopsis thaliana. PLoS One 2011, 6:e26231.

12. Smyth DR, Bowman JL, Meyerowitz EM: Early flower development in Arabidopsis. The Plant Cell Online 1990, 2: 755–767.

13. Yadav RK, Girke T, Pasala S, Xie M, Reddy GV: Gene expression map of the Arabidopsis shoot apical meristem stem cell niche. Proceedings of the National Academy of Sciences 2009, 106: 4941–4946.

14. Yadav RK, Tavakkoli M, Xie M, Girke T, Reddy GV: A high-resolution gene expression map of the Arabidopsis shoot meristem stem cell niche. Development 2014, 141: 2735–2744.

15. Birnbaum K, Shasha DE, Wang JY, Jung JW, Lambert GM, Galbraith DW, Benfey PN: A gene expression map of the Arabidopsis root. Science 2003, 302: 1956–1960.

16. Birnbaum K, Jung JW, Wang JY, Lambert GM, Hirst JA, Galbraith DW, Benfey PN: Cell type–specific expression profiling in plants via cell sorting of protoplasts from fluorescent reporter lines. Nat Methods 2005, 2: 615–619.

17. Carter AD, Bonyadi R, Gifford ML: The use of fluorescence-activated cell sorting in studying plant development and environmental responses. Int J Dev Biol 2013, 57: 545–552.

18. Lan P, Li W, Lin W-D, Santi S, Schmidt W: Mapping gene activity of Arabidopsis root hairs. Genome Biol 2013, 14:R67.

19. Adrian J, Chang J, Ballenger CE, Bargmann BOR, Alassimone J, Davies KA, Lau OS, Matos JL, Hachez C, Lanctot A, Vatén A, Birnbaum KD, Bergmann DC: Transcriptome dynamics of the stomatal lineage: birth, amplification, and termination of a self-renewing population. Dev Cell 2015, 33: 107–118.

20. Ma H, Yanofsky MF, Meyerowitz EM: AGL1-AGL6, an Arabidopsis gene family with similarity to floral homeotic and transcription factor genes. Genes Dev 1991, 5 :484–495.

21. Savidge B, Rounsley SD, Yanofsky MF: Temporal relationship between the transcription of two Arabidopsis MADS box genes and the floral organ identity genes. The Plant Cell Online 1995, 7: 721–733.

22. Colombo M, Brambilla V, Marcheselli R, Caporali E, Kater MM, Colombo L: A new role for the SHATTERPROOF genes during Arabidopsis gynoecium development. Dev Biol 2010, 337: 294–302.

23. Larsson E, Roberts CJ, Claes AR, Franks RG, Sundberg E: Polar Auxin Transport Is Essential for Medial versus Lateral Tissue Specification and Vascular-Mediated Valve Outgrowth in Arabidopsis Gynoecia. Plant Physiol 2014, 166: 1998–2012.

24. Romanel EAC, Schrago CG, Couñago RM, Russo CAM, Alves-Ferreira M: Evolution of the B3 DNA Binding Superfamily: New Insights into REM Family Gene Diversification. PLoS One 2009, 4:e5791.

25. Swaminathan K, Peterson K, Jack T: The plant B3 superfamily. Trends Plant Sci 2008, 13: 647–655.

26. Galbiati F, Sinha Roy D, Simonini S, Cucinotta M, Ceccato L, Cuesta C, Simaskova M, Benkova E, Kamiuchi Y, Aida M, Weijers D, Simon R, Masiero S, Colombo L: An integrative model of the control of ovule primordia formation. Plant J 2013, 76: 446–455.

27. Liljegren SJ, Ditta GS, Eshed Y, Savidge B, Bowman JL, Yanofsky MF: SHATTERPROOF MADS-box genes control seed dispersal in Arabidopsis. Nature 2000, 404: 766–770.

28. Favaro R, Pinyopich A, Battaglia R, Kooiker M, Borghi L, Ditta G, Yanofsky MF, Kater MM, Colombo L: MADS-Box Protein Complexes Control Carpel and Ovule Development in Arabidopsis. Plant Cell2003, 15: 2603–2611.

29. Pinyopich A, Ditta GS, Savidge B, Liljegren SJ, Baumann E, Wisman E, Yanofsky MF: Assessing the redundancy of MADS-box genes during carpel and ovule development. Nature 2003, 424: 85–88.

30. Ó’Maoiléidigh DS, Wellmer F: A Floral Induction System for the Study of Early Arabidopsis Flower Development. In Flower Development. Volume 1110. Edited by Riechmann JL, Wellmer F. New York, NY: Springer New York; 2014: 307–314.

31. Wellmer F, Alves-Ferreira M, Dubois A, Riechmann JL, Meyerowitz EM: Genome-Wide Analysis of Gene Expression during Early Arabidopsis Flower Development. PLoS Genet 2006, 2:e117.

32. Alvarez JP, Goldshmidt A, Efroni I, Bowman JL, Eshed Y: The NGATHA distal organ development genes are essential for style specification in Arabidopsis. Plant Cell 2009, 21: 1373–1393.

33. Trigueros M, Navarrete-Gomez M, Sato S, Christensen SK, Pelaz S, Weigel D, Yanofsky MF, Ferrandiz C: The NGATHA Genes Direct Style Development in the Arabidopsis Gynoecium. THE PLANT CELL ONLINE 2009, 21: 1394–1409.

34. Lamesch P, Berardini TZ, Li D, Swarbreck D, Wilks C, Sasidharan R, Muller R, Dreher K, Alexander DL, Garcia-Hernandez M, Karthikeyan AS, Lee CH, Nelson WD, Ploetz L, Singh S, Wensel A, Huala E: The Arabidopsis Information Resource (TAIR): improved gene annotation and new tools. Nucleic Acids Res 2012, 40 (Database issue):D1202–10.

35. Trapnell C, Roberts A, Goff L, Pertea G, Kim D, Kelley DR, Pimentel H, Salzberg SL, Rinn JL, Pachter L: Differential gene and transcript expression analysis of RNA-seq experiments with TopHat and Cufflinks. Nat Protoc 2012, 7: 562–578.

36. Robinson MD, McCarthy DJ, Smyth GK: edgeR: a Bioconductor package for differential expression analysis of digital gene expression data. Bioinformatics 2010, 26 :139–140.

37. Love MI, Huber W, Anders S: Moderated estimation of fold change and dispersion for RNA-seq data with DESeq2. Genome Biol 2014, 15.

38. Gremski K, Ditta G, Yanofsky MF: The HECATE genes regulate female reproductive tract development in Arabidopsis thaliana. Development 2007, 134: 3593–3601.

39. Kuusk S, Sohlberg JJ, Long JA, Fridborg I, Sundberg E: STY1 and STY2 promote the formation of apical tissues during Arabidopsis gynoecium development. Development 2002, 129: 4707–4717.

40. Skinner DJ, Gasser CS: Expression-based discovery of candidate ovule development regulators through transcriptional profiling of ovule mutants. BMC Plant Biol 2009, 9: 29.

41. Wuest SE, Vijverberg K, Schmidt A, Weiss M, Gheyselinck J, Lohr M, Wellmer F, Rahnenführer J, Mering C von, Grossniklaus U: Arabidopsis female gametophyte gene expression map reveals similarities between plant and animal gametes. Curr Biol 2010, 20:506-512.

42. Bowman JL, Smyth DR: CRABS CLAW, a gene that regulates carpel and nectary development in Arabidopsis, encodes a novel protein with zinc finger and helixloop-helix domains. Development 1999, 126:2387–96.

43. Azhakanandam S, Nole-Wilson S, Bao F, Franks RG: SEUSS and AINTEGUMENTA Mediate Patterning and Ovule Initiation during Gynoecium Medial Domain Development. Plant Physiol 2008, 146:1165–1181.

44. Heisler M, Atkinson A, Bylstra YH, Walsh R, Smyth DR: SPATULA, a gene that controls development of carpel margin tissues in Arabidopsis, encodes a bHLH protein. Development 2001, 128: 1089–1098.

45. Kamiuchi Y, Yamamoto K, Furutani M, Tasaka M, Aida M: The CUC1 and CUC2 genes promote carpel margin meristem formation during Arabidopsis gynoecium development. Front Plant Sci 2014, 5.

46. Bennett MJ, Marchant A, Green HG, May ST, Ward SP, Millner PA, Walker AR, Schulz B, Feldmann KA: Arabidopsis AUX1 gene: a permease-like regulator of root gravitropism. Science 1996, 273: 948–950.

47. Cheng Y, Dai X, Zhao Y: Auxin biosynthesis by the YUCCA flavin monooxygenases controls the formation of floral organs and vascular tissues in Arabidopsis. Genes Dev 2006, 20: 1790–1799.

48. Pagnussat GC, Yu H-J, Ngo QA, Rajani S, Mayalagu S, Johnson CS, Capron A, Xie L-F, Ye D, Sundaresan V: Genetic and molecular identification of genes required for female gametophyte development and function in Arabidopsis. Development 2005, 132:603-614.

49. Payne CT, Zhang F, Lloyd AM: GL3 encodes a bHLH protein that regulates trichome development in arabidopsis through interaction with GL1 and TTG1. Genetics 2000, 156:1349–1362.

50. Crawford BCW, Yanofsky MF: HALF FILLED promotes reproductive tract development and fertilization efficiency in Arabidopsis thaliana. Development 2011, 138: 2999–3009.

51. Transcription Factor Enrichment Calculator [https://dgrinevich.shinyapps.io/ShinyTF]

52. Franco-Zorrilla JM, Cubas P, Jarillo JA, Fernandez-Calvin B, Salinas J, Martinez-Zapater JM: AtREM1, a Member of a New Family of B3 Domain-Containing Genes, Is Preferentially Expressed in Reproductive Meristems. Plant Physiol 2002, 128: 418–427.

53. Mantegazza O, Gregis V, Mendes MA, Morandini P, Alves-Ferreira M, Patreze CM, Nardeli SM, Kater MM, Colombo L: Analysis of the arabidopsis REM gene family predicts functions during flower development. Ann Bot 2014, 114: 1507–1515.

54. Mantegazza O, Gregis V, Chiara M, Selva C, Leo G, Horner DS, Kater MM: Gene coexpression patterns during early development of the native Arabidopsis reproductive meristem: novel candidate developmental regulators and patterns of functional redundancy. Plant J2014, 79: 861–877.

55. Matias-Hernandez L, Battaglia R, Galbiati F, Rubes M, Eichenberger C, Grossniklaus U, Kater MM, Colombo L: VERDANDI Is a Direct Target of the MADS Domain Ovule Identity Complex and Affects Embryo Sac Differentiation in Arabidopsis. Plant Cell 2010, 22: 1702–1715.

56. Mizzotti C, Ezquer I, Paolo D, Rueda-Romero P, Guerra RF, Battaglia R, Rogachev I, Aharoni A, Kater MM, Caporali E, Colombo L: SEEDSTICK is a Master Regulator of Development and Metabolism in the Arabidopsis Seed Coat. PLoS Genet 2014, 10:e1004856.

57. Phillips AL, Ward DA, Uknes S, Appleford NE, Lange T, Huttly AK, Gaskin P, Graebe JE, Hedden P: Isolation and expression of three gibberellin 20-oxidase cDNA clones from Arabidopsis. Plant Physiol 1995, 108: 1049–1057.

58. Jasinski S, Piazza P, Craft J, Hay A, Woolley L, Rieu I, Phillips A, Hedden P, Tsiantis M: KNOX Action in Arabidopsis Is Mediated by Coordinate Regulation of Cytokinin and Gibberellin Activities. Curr Biol 2005, 15: 1560–1565.

59. Hay A, Angela H, Hardip K, Andrew P, Peter H, Sarah H, Miltos T: The Gibberellin Pathway Mediates KNOTTED1-Type Homeobox Function in Plants with Different Body Plans. Curr Biol 2002, 12: 1557–1565.

60. Guida A, Lindstädt C, Maguire SL, Ding C, Higgins DG, Corton NJ, Berriman M, Butler G: Using RNA-seq to determine the transcriptional landscape and the hypoxic response of the pathogenic yeast Candida parapsilosis. BMC Genomics 2011, 12: 628.

61. Bradford JR, Hey Y, Yates T, Li Y, Pepper SD, Miller CJ: A comparison of massively parallel nucleotide sequencing with oligonucleotide microarrays for global transcription profiling. BMC Genomics 2010, 11: 282.

62. Mudge J, Miller NA, Khrebtukova I, Lindquist IE, May GD, Huntley JJ, Luo S, Zhang L, van Velkinburgh JC, Farmer AD, Lewis S, Beavis WD, Schilkey FD, Virk SM, Black CF, Myers MK, Mader LC, Langley RJ, Utsey JP, Kim RW, Roberts RC, Khalsa SK, Garcia M, Ambriz-Griffith V, Harlan R, Czika W, Martin S, Wolfinger RD, Perrone-Bizzozero NI, Schroth GP, et al.: Genomic Convergence Analysis of Schizophrenia: mRNA Sequencing Reveals Altered Synaptic Vesicular Transport in Post-Mortem Cerebellum. PLoS One 2008, 3:e3625.

63. Nookaew I, Papini M, Pornputtapong N, Scalcinati G, Fagerberg L, Uhlen M, Nielsen J: A comprehensive comparison of RNA-Seq-based transcriptome analysis from reads to differential gene expression and cross-comparison with microarrays: a case study in Saccharomyces cerevisiae. Nucleic Acids Res 2012, 40: 10084–10097.

64. Wang C, Gong B, Bushel PR, Thierry-Mieg J, Thierry-Mieg D, Xu J, Fang H, Hong H, Shen J, Su Z, Meehan J, Li X, Yang L, Li H, Łabaj PP, Kreil DP, Megherbi D, Gaj S, Caiment F, van Delft J, Kleinjans J, Scherer A, Devanarayan V, Wang J, Yang Y, Qian H-R, Lancashire LJ, Bessarabova M, Nikolsky Y, Furlanello C, et al.: The concordance between RNA-seq and microarray data depends on chemical treatment and transcript abundance. Nat Biotechnol 2014, 32: 926–932.

65. Xu X, Zhang Y, Williams J, Antoniou E, McCombie WR, Wu S, Zhu W, Davidson NO, Denoya P, Li E: Parallel comparison of Illumina RNA-Seq and Affymetrix microarray platforms on transcriptomic profiles generated from 5-aza-deoxy-cytidine treated HT-29 colon cancer cells and simulated datasets. BMC Bioinformatics 2013, 14(Suppl 9):S1.

66. Zhao S, Fung-Leung W-P, Bittner A, Ngo K, Liu X: Comparison of RNA-Seq and Microarray in Transcriptome Profiling of Activated T Cells. PLoS One 2014, 9:e78644.

67. Marioni JC, Mason CE, Mane SM, Stephens M, Gilad Y: RNA-seq: An assessment of technical reproducibility and comparison with gene expression arrays. Genome Res 2008, 18: 1509–1517.

68. Halbritter F, Vaidya HJ, Tomlinson SR: GeneProf: analysis of high-throughput sequencing experiments. Nat Methods 2012, 9: 7–8.

69. Sims D, Sudbery I, Ilott NE, Heger A, Ponting CP: Sequencing depth and coverage: key considerations in genomic analyses. Nat Rev Genet 2014, 15: 121–132.

70. Franks RG, Wang C, Levin JZ, Liu Z: SEUSS, a member of a novel family of plant regulatory proteins, represses floral homeotic gene expression with LEUNIG. Development 2002, 129: 253–263.

71. Ayoubi TA, Van De Ven WJ: Regulation of gene expression by alternative promoters. The FASEB Journal 1996, 10: 453–460.

72. Drakakaki G, Zabotina O, Delgado I, Robert S, Keegstra K, Raikhel N: Arabidopsis Reversibly Glycosylated Polypeptides 1 and 2 Are Essential for Pollen Development. Plant Physiol 2006, 142: 1480–1492.

73. Sauer M, Robert S, Kleine-Vehn J: Auxin: simply complicated. J Exp Bot 2013, 64: 2565–2577.

74. Woodward AW, Bartel B: Auxin: regulation, action, and interaction. Ann Bot 2005, 95: 707–735.

75. Sehra B, Franks RG: Auxin and cytokinin act during gynoecial patterning and the development of ovules from the meristematic medial domain. Wiley Interdiscip Rev Dev Biol 2015.

76. Moubayidin L, Ostergaard L: Dynamic Control of Auxin Distribution Imposes a Bilateral-to-Radial Symmetry Switch during Gynoecium Development. Curr Biol 2014, 24: 2743–2748.

77. Stepanova AN, Robertson-Hoyt J, Yun J, Benavente LM, Xie D-Y, Doležal K, Schlereth A, Jürgens G, Alonso JM: TAA1-Mediated Auxin Biosynthesis Is Essential for Hormone Crosstalk and Plant Development. Cell 2008, 133: 177–191.

78. Tao Y, Ferrer JL, Ljung K, Pojer F, Hong F, Long JA, Li L, Moreno JE, Bowman ME, Ivans LJ, Cheng Y, Lim J, Zhao Y, Ballare CL, Sandberg G, Noel JP, Chory J: Rapid synthesis of auxin via a new tryptophan-dependent pathway is required for shade avoidance in plants. Cell 2008, 133: 164–176.

79. Zhao Y, Christensen SK, Fankhauser C, Cashman JR, Cohen JD, Weigel D, Chory J: A role for flavin monooxygenase-like enzymes in auxin biosynthesis. Science 2001, 291: 306–309.

80. Martinez-Fernandez I, Sanchis S, Marini N, Balanza V, Ballester P, Navarrete-Gomez M, Oliveira AC, Colombo L, Ferrandiz C: The effect of NGATHA altered activity on auxin signaling pathways within the Arabidopsis gynoecium. Front Plant Sci 2014, 5: 210.

81. Benkova E, Michniewicz M, Sauer M, Teichmann T, Seifertova D, Jurgens G, Friml J: Local, efflux-dependent auxin gradients as a common module for plant organ formation. Cell 2003, 115: 591–602.

82. Blilou I, Xu J, Wildwater M, Willemsen V, Paponov I, Friml J, Heidstra R, Aida M, Palme K, Scheres B: The PIN auxin efflux facilitator network controls growth and patterning in Arabidopsis roots. Nature 2005, 433: 39–44.

83. Wu M-F, Tian Q, Reed JW: Arabidopsis microRNA167 controls patterns of ARF6 and ARF8 expression, and regulates both female and male reproduction. Development 2006, 133: 4211–4218.

84. Nagpal P, Ellis CM, Weber H, Ploense SE, Barkawi LS, Guilfoyle TJ, Hagen G, Alonso JM, Cohen JD, Farmer EE, Ecker JR, Reed JW: Auxin response factors ARF6 and ARF8 promote jasmonic acid production and flower maturation. Development 2005,132:4107–4118.

85. Williams L, Carles CC, Osmont KS, Fletcher JC: A database analysis method identifies an endogenous trans-acting short-interfering RNA that targets the Arabidopsis ARF2, ARF3, and ARF4 genes. Proc Natl Acad Sci USA 2005, 102: 9703–9708.

86. Wang J-W, Schwab R, Czech B, Mica E, Weigel D: Dual effects of miR156-targeted SPL genes and CYP78A5/KLUH on plastochron length and organ size in Arabidopsis thaliana. Plant Cell 2008, 20: 1231–1243.

87. Wu G, Poethig RS: Temporal regulation of shoot development in Arabidopsis thaliana by miR156 and its target SPL3. Development 2006, 133: 3539–3547.

88. Zhang T-Q, Lian H, Tang H, Dolezal K, Zhou C-M, Yu S, Chen J-H, Chen Q, Liu H, Ljung K, Wang J-W: An intrinsic microRNA timer regulates progressive decline in shoot regenerative capacity in plants. Plant Cell 2015, 27: 349–360.

89. Wang J-W, Czech B, Weigel D: miR156-regulated SPL transcription factors define an endogenous flowering pathway in Arabidopsis thaliana. Cell 2009, 138: 738–749.

90. Leal Valentim F, Sv M, Posé D, Kim MC, Schmid M, van Ham R: A Quantitative and Dynamic Model of the Arabidopsis Flowering Time Gene Regulatory Network. PLoS One 2015, 10:e0116973.

91. Gandikota M, Birkenbihl RP, Höhmann S, Cardon GH, Saedler H, Huijser P: The miRNA156/157 recognition element in the 3′ UTR of the Arabidopsis SBP box gene SPL3 prevents early flowering by translational inhibition in seedlings. Plant J 2007, 49: 683–693.

92. Heberle H, Meirelles GV, da Silva FR, Telles GP, Minghim R: InteractiVenn: a web-based tool for the analysis of sets through Venn diagrams. BMC Bioinformatics 2015, 16.

93. [http://www.interactivenn.net] webcite

94. Moussaieff A, Rogachev I, Brodsky L, Malitsky S, Toal TW, Belcher H, Yativ M, Brady SM, Benfey PN, Aharoni A: High-resolution metabolic mapping of cell types in plant roots. Proceedings of the National Academy of Sciences 2013.

95. Petricka JJ, Schauer MA, Megraw M, Breakfield NW, Thompson JW, Georgiev S, Soderblom EJ, Ohler U, Moseley MA, Grossniklaus U, Benfey PN: The protein expression landscape of the Arabidopsis root. Proc Natl Acad Sci US A 2012, 109: 6811–6818.

96. Li S, Liberman LM, Mukherjee N, Benfey PN, Ohler U: Integrated detection of natural antisense transcripts using strand-specific RNA sequencing data. Genome Res 2013, 23: 1730–1739.

97. Breakfield NW, Corcoran DL, Petricka JJ, Shen J, Sae-Seaw J, Rubio-Somoza I, Weigel D, Ohler U, Benfey PN: High-resolution experimental and computational profiling of tissue-specific known and novel miRNAs in Arabidopsis. Genome Res 2012, 22: 163–176.

98. Petersson SV, Johansson AI, Kowalczyk M, Makoveychuk A, Wang JY, Moritz T, Grebe M, Benfey PN, Sandberg G, Ljung K: An auxin gradient and maximum in the Arabidopsis root apex shown by high-resolution cell-specific analysis of IAA distribution and synthesis. Plant Cell 2009, 21: 1659–1668.

99. Earley KW, Haag JR, Pontes O, Opper K, Juehne T, Song K, Pikaard CS: Gateway-compatible vectors for plant functional genomics and proteomics. Plant J 2006, 45: 616–629.

100. Nakagawa T, Kurose T, Hino T, Tanaka K, Kawamukai M, Niwa Y, Toyooka K, Matsuoka K, Jinbo T, Kimura T: Development of series of gateway binary vectors, pGWBs, for realizing efficient construction of fusion genes for plant transformation. J Biosci Bioeng 2007, 104: 34–41.

101. Schmittgen TD, Livak KJ: Analyzing real-time PCR data by the comparative CT method. Nat Protoc 2008, 3: 1101–1108.

102. [http://sourceforge.net/projects/yjlee-r-packages/files/bear/] webcite

103. [https://cran.r-project.org/web/packages/plyr/index.html] webcite

104. [https://cran.r-project.org/web/packages/ggplot2/index.html] webcite

105. Schmieder R, Edwards R: Quality control and preprocessing of metagenomic datasets. Bioinformatics 2011, 27: 863–864.

106. Lindgreen S: AdapterRemoval: easy cleaning of next-generation sequencing reads. BMC Res Notes 2012, 5: 337.

107. Trapnell C, Pachter L, Salzberg SL: TopHat: discovering splice junctions with RNA-Seq. Bioinformatics 2009, 25: 1105–1111.

108. [http://support.illumina.com/sequencing/sequencing_software/igenome.html] webcite

109. Li H, Handsaker B, Wysoker A, Fennell T, Ruan J, Homer N, Marth G, Abecasis G, Durbin R, 1000 Genome Project Data Processing Subgroup: The Sequence Alignment/Map format and SAMtools. Bioinformatics 2009, 25: 2078–2079.

110. Anders S, Pyl PT, Huber W: HTSeq-a Python framework to work with high-throughput sequencing data. Bioinformatics 2015, 31: 166–169.

111. Anders S, Huber W: Differential expression analysis for sequence count data. Genome Biol 2010, 11:R106.

112. Rau A, Gallopin M, Celeux G, Jaffrezic F: Data-based filtering for replicated high-throughput transcriptome sequencing experiments. Bioinformatics 2013, 29: 2146–2152.

113. Soneson C, Delorenzi M: A comparison of methods for differential expression analysis of RNA-seq data. BMC Bioinformatics 2013, 14: 91.

114. Bourgon R, Gentleman R, Huber W: Independent filtering increases detection power for high-throughput experiments. Proceedings of the National Academy of Sciences 2010, 107: 9546–9551.

115. Pimentel H, Parra M, Gee S, Ghanem D, An X, Li J, Mohandas N, Pachter L, Conboy JG: A dynamic alternative splicing program regulates gene expression during terminal erythropoiesis. Nucleic Acids Res 2014, 42: 4031–4042.

116. Seyednasrollah F, Laiho A, Elo LL: Comparison of software packages for detecting differential expression in RNA-seq studies. Brief Bioinform 2015, 16: 59–70.

117. Suzuki A, Matsushima K, Makinoshima H, Sugano S, Kohno T, Tsuchihara K, Suzuki Y: Single-cell analysis of lung adenocarcinoma cell lines reveals diverse expression patterns of individual cells invoked by a molecular target drug treatment. Genome Biol 2015, 16.

118. Yilmaz A, Mejia-Guerra MK, Kurz K, Liang X, Welch L, Grotewold E: AGRIS: the Arabidopsis Gene Regulatory Information Server, an update. Nucleic Acids Res 2011, 39 (Database):D1118-D1122.

119. Chen H, Boutros PC: VennDiagram: a package for the generation of highly-customizable Venn and Euler diagrams in R. BMC Bioinformatics 2011, 12: 35.

120. [http://cran.r-project.org/web/packages/pheatmap/index.html] webcite

121. [http://stat.ethz.ch/R-manual/R-devel/library/grDevices/html/colorRamp.html] webcite

122. Alexa A, Rahnenführer J: topGO: Enrichment Analysis for Gene Ontology; 2010. Bioconductor Package Version. 2009.

123. Maere S, Heymans K, Kuiper M: BiNGO: a Cytoscape plugin to assess overrepresentation of Gene Ontology categories in Biological Networks. Bioinformatics 2005, 21: 3448–3449.

124. Shannon P, Markiel A, Ozier O, Baliga NS, Wang JT, Ramage D, Amin N, Schwikowski B, Ideker T: Cytoscape: a software environment for integrated models of biomolecular interaction networks. Genome Res 2003, 13: 2498–2504.

125. Schneitz K, Hülskamp M, Pruitt RE: Wild-type ovule development in Arabidopsis thaliana: a light microscope study of cleared whole-mount tissue. Plant J 1995, 7: 731–749.

126. Ashburner M, Ball CA, Blake JA, Botstein D, Butler H, Cherry JM, Davis AP, Dolinski K, Dwight SS, Eppig JT, Harris MA, Hill DP, Issel-Tarver L, Kasarskis A, Lewis S, Matese JC, Richardson JE, Ringwald M, Rubin GM, Sherlock G: Gene ontology: tool for the unification of biology. The Gene Ontology Consortium. Nat Genet 2000, 25: 25–29.

